# On the evolution, function and cellular fate of *Neurospora crassa* ACW-1 and NCW-3, proteins with different cell wall interaction mechanism

**DOI:** 10.64898/2026.05.09.718313

**Authors:** Ana Sofía Ramírez-Pelayo, Olga A. Callejas-Negrete, Lorena Amaya-Delgado, Jorge Verdín

## Abstract

The fungal cell wall is populated by proteins (CWPs), mostly uncharacterized, that show an atypical evolutionary behavior. Most CWPs are glycosylphosphatidylinositol(GPI)-proteins, followed by proteins with internal repeats (PIR), and non-covalently attached proteins that harbor carbohydrate binding domains (CBM). Several structural CWPs are initially bound to the same wall carbohydrates, but either covalently or non-covalently. However, it is not clear whether they work in the same way and if they are subjected to the same evolutionary constraints. In *Neurospora crassa*, CWPs ACW-1 (NCU08936) and NCW-3 (NCU07817) bind to β-1,3-glucans through a GPI anchor or a predicted CBM-52 domain, respectively. Here, the evolutionary trajectories and functional roles of both CWPs were analyzed. Both proteins localized primarily to distal septa and hyphal wall surfaces. Morphological characterization and stress cell wall assays suggested that both proteins contribute to cell wall integrity, but NCW-3 likely plays a more prominent role. ACW-1 and NCW-3 homologues were predominantly identified in Ascomycota. ACW-1 displayed a broader distribution than NCW-3, whose homologues were largely restricted to Sordariales. Despite these differences, both protein families exhibited similar moderate global conservation and signatures of purifying selection within shared taxa. Nevertheless, a divergence gradient was identified within ACW-1, related to its tandem leucine-rich repeat (LRR) regions. A similar local accumulation of evolutionary change was not observed within NCW-3. These findings suggested that distinct CWP architectures can accommodate different patterns of sequence diversification despite sharing similar global evolutionary change.

## INTRODUCTION

Fungal cells are surrounded by a cell wall that shapes their morphogenesis, growth and survival. It provides mechanical support and mediates interactions with the extracellular environment [1]. The cell wall is generally built of cross-linked glucans (β-1,3-, β-1,3/β-1,4-, β-1,6- and/or α-1,3-glucans), chitin, chitosan, pigments and cell wall resident glycoproteins (CWPs). Recent solid-state nuclear magnetic resonance (NMR) assays [2–4] of intact *Aspergillus fumigatus* cell walls suggest these components are organized in an inner domain and a highly mobile outer shell. The inner domain is divided into a hydrated mobile layer of β-glucans (β-1,6-, β-1,4- and β-1,3-glucans) and a hydrophobic, rigid scaffold of cross-linked α-1,3-glucan and chitin. The outer highly mobile layer is composed mainly of α-1,3-glucans, galactomannan and CWPs.

The cell wall is a highly dynamic structure whose composition varies with species, growth stage and environmental conditions. This versatility enables fungi to withstand a wide diversity of stresses, including the presence of antifungal drugs [5,6], loss of major cell wall polysaccharides (β-1,3-glucans, chitin and mannan), and changes in the extracellular environment [3]. The fungal cell wall dynamic composition and its ability to respond to stimuli involve multiple proteins, including cytosolic factors, membrane-bound and CWPs.

CWPs play key roles in modifying cell wall polysaccharides, transducing extracellular signals and facilitating adhesion and surface invasion. CWPs are typically post-translationally modified with N- and/or O-glycosylations [7,8] and can be incorporated into the wall covalently or noncovalently. In yeasts, most covalently attached CWPs are glycosylphosphatidylinositol (GPI)-anchored, which integrate into the cell wall through a transglycosylation reaction between the processed GPI glycan remnant and the β-1,6-glucan of the cell wall [9]. Proteins with internal repeats (PIR), the second most abundant class of CWPs, integrate covalently either via an alkali-sensitive ester linkage between a glutamine (Q) residue of their PIR motifs and β-1,3-glucans, or through disulfide bonds between their C-terminal cysteine (Cys)-rich domains and other CWPs [10,11]. Non-covalently attached CWPs interact with the cell wall through electrostatic and hydrophobic interactions, as well as carbohydrate binding modules (CBM) [12,13]. GPI- and non-covalently attached CWPs can be additionally cross-linked to the wall through their mannosylations to cell wall polysaccharides [14].

Many fungal CWPs amino acid sequences are enriched in low-complexity regions and tandemly repeated sequences: adhesins, flocculins, virulence factors, PIR proteins, structural CWPs and proteins associated with cell wall remodelling and stress response [15–17]. Comparative genomic analyses have shown that CWPs frequently display high sequence variability and lineage specificity [17–19], which has been interpreted as evidence of rapid evolutionary change relative to more conserved cytosolic proteins [17,19]. Tandem repeat regions have been proposed as an expression of this variability, as the repetitive sequences makes them prone to expansion and/or contraction through processes such as DNA replication slippage, unequal cross-over, or double strand break repair [20–22]. Among PIR-proteins, whole protein polarity inversion, as well as generation of inverted sequence (mirror-like) PIR motifs have been identified [23]. These changes can alter repeat number and sequence variability, potentially contributing to functional diversification [18,21]. In *Neurospora crassa*, more than 40 CWPs have been described. These include glycosyl hydrolases involved in the formation of the cell wall, like GH16 glucanosyltransferases [24–26], GH17 glucanosylhydrolases/transferases [26,27], GH72 glucanosyltransferases [14,28–30], GH76 mannosyltransferases [29] and many other uncharacterized CWPs [31]. Among these, ACW-1 (anchored cell wall protein 1, NCU08936), also annotated as CCG-15 (clock-controlled protein 15), and NCW-3 (non-anchored cell wall protein 3, NCU07817) have been described as integral CWPs present in vegetative cells and conidial cell walls. ACW-1 is homologous to *S. cerevisiae* Ecm33p, which is involved in the maintenance of cell wall integrity [32]. ACW-1 is a GPI-protein although there is evidence that both the presence of a GPI addition machinery [31,33] and the processing and cross-linking of its N-glycosylation to β-1,3/1,4-glucan [14,29,34,35] are needed for the stable incorporation and anchoring into the cell wall. NCW-3 has no known characterized homologues. It contains a predicted N-terminal CAZy family 52 carbohydrate binding module (CBM-52), which in fission yeast *Schizosaccharomyces pombe* binds to the linear, insoluble β-1,3-glucan found in the primary septum [36]. NCW-3 has been proposed to be covalently bound to the cell wall through O-glycosylation processing [31]. It is necessary to produce mature perithecia [37], and it is upregulated in *N. crassa* Δ*gul-1* cells and non-viscous liquid cultures [38]. GUL-1 is part of the COT-1 pathway and is involved in multiple cellular processes, including the regulation of polarized hyphal growth and cell wall remodelling [39].

Within this framework, ACW-1 and NCW-3 are non-catalytic, structural, cell wall proteins that are ultimately recovered in the covalently associated cell wall fraction, but appear to differ in their mechanisms of initial association with β-1,3-glucans. ACW-1 has a GPI anchor, which is known to mediate targeting and subsequent integration of proteins into the fungal cell wall. In contrast, NCW-3 contains a predicted CBM-52 domain, which is expected to mediate specific non-covalent recognition of this polysaccharide within the cell wall matrix [30]. In both cases, these domain-mediated interactions likely precede and enable posttranslational covalent incorporation processes, such as GPI- or N-/O-glycosylation-dependent cross-linking, which contribute to stable retention within the wall.

These structural differences suggest that ACW-1 and NCW-3 operate through distinct cell wall targeting modules that may impose distinct functional constraints. Such differences could be associated with divergent evolutionary patterns among evolutionarily atypical CWPs. By integrating evolutionary, structural and functional analyses, this work examines whether ACW-1 and NCW-3 follow similar evolutionary trajectories or display distinct patterns of sequence evolution within *N. crassa*.

## RESULTS

### ACW-1 and NCW-3 homologues were predominantly identified in Ascomycota with diverse lifestyles

A Fungal Tree of Life (FtoL) of 4147 fungal species was reconstructed from genome-scale data from NCBI RefSeq database [40] (September 2024) to establish a species-level phylogenetic framework (**Figure 1**). This reconstruction was used to assess the distribution of ACW-1 and NCW-3 homologues across fungal lineages and explore its evolutionary history. A total of 1136 ACW-1 homologues were identified predominantly in Ascomycota, with a few in Zoopagomycota (Dimargaritales, Harpellales) and Mucoromycota (Mortierellales) (**Figure 1A**). ACW- 1 homologues were distributed across 76 orders and 179 families (for a comprehensive breakdown, see **Table S1**) with *Aspergillaceae* (Eurotiales) having the highest number of sequences (106), followed by *Nectriaceae* (78, Hypocreales), *Xylariaceae* (51, Xylariales), *Saccharomycetaceae* (45, Saccharomycetales), and Glomerellaceae (45, Glomerellales). These fungi spanned various ecological niches, predominantly plant-associated pathogens and endophytes, saprotrophs, and extremophiles. Two to four ACW-1 paralogs were present in yeasts (*Saccharomycetaceae*, *Pichiaceae*, *Saccharomycodaceae*, *Wickerhamomycetaceae*, *Phaffomycetaceae*, *Debaryomycetaceae*, *Metschnikowaceae*, *Dipodascaceae* and *Pachysolenaceae*), in species with diverse lifestyles such as plant and human pathogens, saprotrophs, methylotrophs, spoilage yeasts, and associated with fermentation. Two ACW-1 paralogs were also found in some endophytic fungi belonging to families *Xylariaceae* and *Hypoxylaceae* (Xylariales), one extremotolerant rock-inhabiting black fungus in *Trichomeriaceae* (Chaetothyriales), a plant pathogen in *Botryosphaeriaceae* (Botryosphaeriales), an extremophilic fungi isolated from rocks and lichens in Antarctica in *Teratosphaeriaceae* (Mycosphaerellales), and a plant pathogen in *Gnomoniaceae* (Diaporthales). This indicated ACW-1 is not restricted to a specific ecological lifestyle.

**Figure 1.**
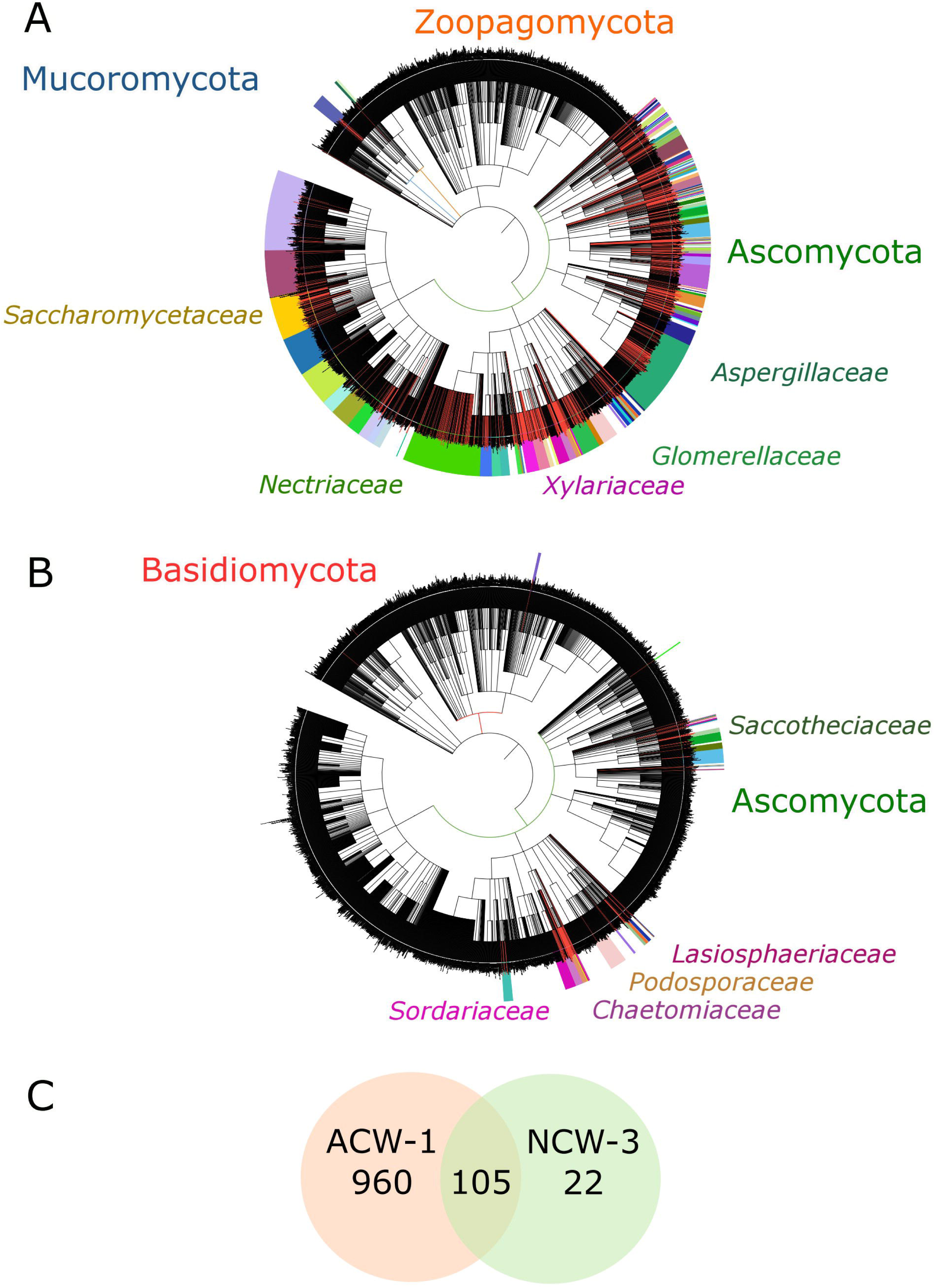
ACW-1 and NCW-3 were majorly restricted to Ascomycota. A) 1136 ACW-1 homologues were majorly found in Ascomycota, distributed in 76 orders across 179 families, of which Aspergillaceae (Eurotiales) was the most numerous with 106 sequences, followed by Nectriaceae (78, Hypocreales), Xylariaceae (51, Xylariales), more than one paralog was found in Saccharomycetaceae (45, Saccharomycetales), and Glomerellaceae (45, Glomerellales); nine homologues were present in Mucoromycota and two in Zoopagomycota. B) 140 NCW-3 homologues were also almost exclusively found in Ascomycota, constrained to Sordariales, in families such as Chaetomiaceae (17), Podosporaceae (16), Lasiophaeriaceae (11) and Sordariaceae (8), and Dothideales in Saccotheciaceae (10); four homologues were found in Basidiomycota. ACW-1 and NCW-3 homologues were searched in 4147 species. In the Fungal Tree of Life (FTOL), colored branches indicate a species that encodes at least one ACW-1 or NCW-3 homologue. The outermost circle groups and labels fungal species in families. Fungal species phylogeny was retrieved from 4147 reference proteomes downloaded from the RefSeq NCBI database [40] (September 2024), and the presence of ACW-1 or NCW-3 in fungal families classes was annotated with iTOL v7.2.2 [116]. C) Only 105 fungal species encode for homologues of both ACW-1 and NCW-3; ACW-1 homologues were found exclusively in 960 species, while only 22 species encode NCW-3 homologues. The Venn diagram was generated in Inkscape [129].

For NCW-3, 139 homologues were also found mostly in Ascomycota (**Figure 1B**). However, in contrast to the broad distribution of ACW-1, NCW-3 homologues were restricted to a narrower taxonomic range, distributed across 21 orders and 37 families (**Table S2**); 4 homologues were identified in Basidiomycota (Agaricales). The most abundant family was *Chaetomiaceae* (17, Sordariales), followed by *Podosporaceae* (16, Sordariales), *Lasiosphaeriaceae* (11, Sordariales), *Saccotheciaceae* (10, Dothideales) and *Sordariaceae* (8, Sordariales). Homologues were found in saprotrophs, coprophilous fungi, endophytes, plant pathogens and some extremophiles. NCW-3 paralogs were also found in 11 fungi almost exclusively in Sordariales (*Lasiosphaeriaceae*, *Chaetomiaceae*, *Podosporaceae*, *Schizotheciaceae*), with one in Pleosporales (*Melanommataceae*) and another in Cephalothecales (*Cephalothecaceae*). ACW-1 and NCW-3 homologues were both simultaneously present in 105 species. ACW-1 homologues mainly occurred by themselves in 960 species, while NCW-3 homologues in only 22 (**Figure 1C**). This asymmetry indicated that ACW-1 is more broadly encoded, whereas NCW-3 may represent a more restricted or lineage-specific component, which is consistent with distinct evolutionary trajectories within Ascomycota.

### ACW-1 and NCW-3 phylogenies showed patterns not strictly congruent with the species-level phylogeny

Maximum likelihood phylogenies constructed from orthologous protein sequences of ACW-1 and NCW-3 were generated to infer gene family evolutionary relationships, which were then compared to the species-level phylogeny to assess congruence (**Figure 2**). ACW-1 homologues segregated into 8 distinct clades that broadly corresponded to taxonomic families (**Figure 2A**). However, the ACW-1 clades distribution showed a topology that deviated from the species-level phylogeny (**Figure 1A**). *Glomerellaceae*, *Xylariaceae* and *Nectriaceae* were grouped under the same clade in the species-level phylogeny (**Figure 1A**); however, in the ACW-1-based phylogeny, only the homologues of *Glomerellaceae* and *Xylariaceae* were grouped within the same clade (**Figure 2A**). *Aspergillaceae* sequences were placed in a more distant clade relative to the species-level phylogeny. In the ACW-based phylogeny, the farthest branches corresponded to Saccharomycotina yeasts (*Saccharomycetaceae*, *Saccharomycodaceae*, *Debaryomycetaceae*, *Pachysolanceae*, *Pichiaceae*, *Wickerhammomycetaceae*), which may reflect lineage-specific diversification patterns. *Saccharomycetaceae* yeasts further split into two distinct subclades, each including paralogs.

**Figure 2.**
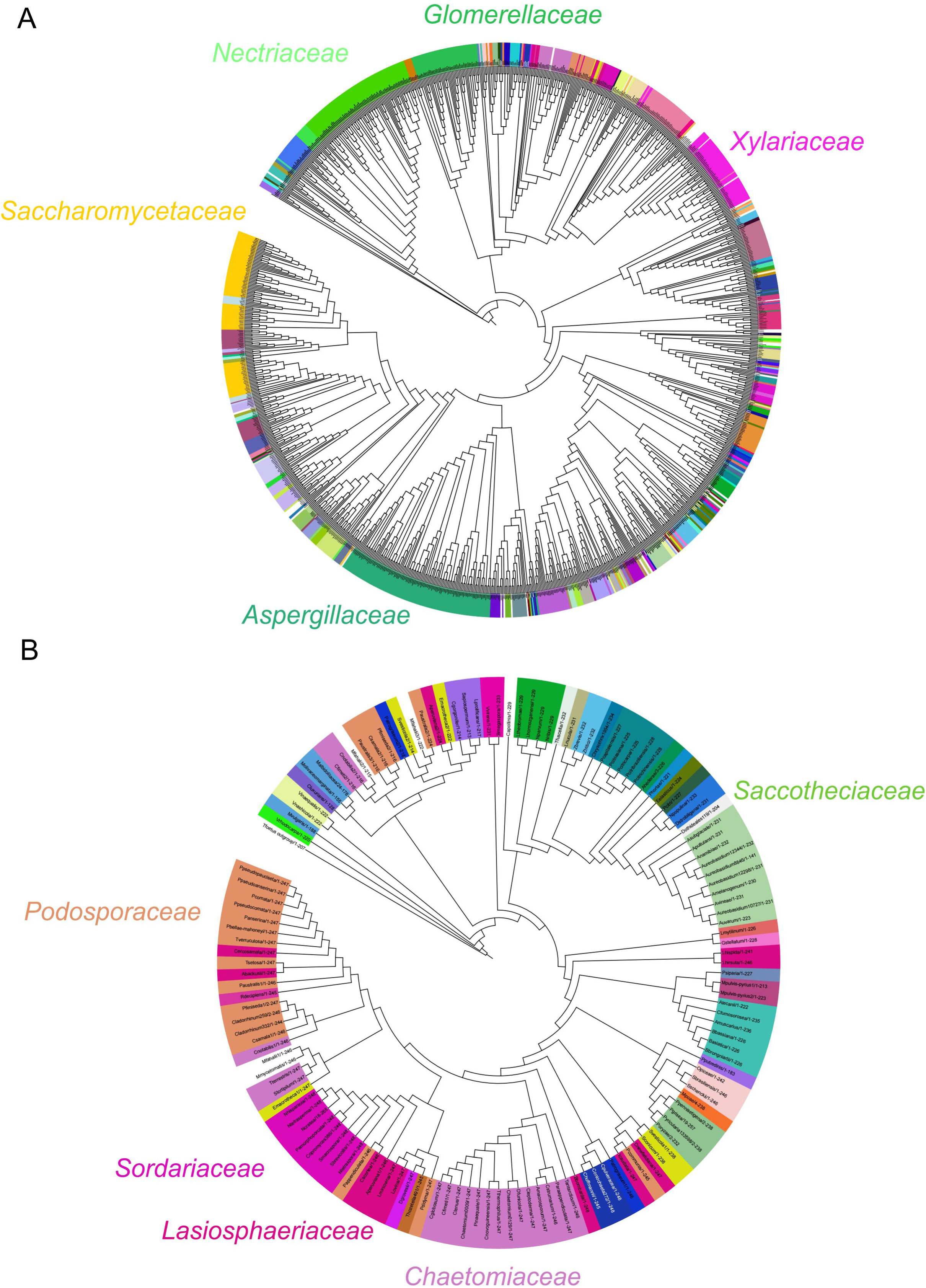
Distribution of ACW-1 and NCW-3 homologues in phylogenetic trees. A) Maximum-likelihood phylogenetic reconstruction of ACW-1 homologues showed their segregation into eight distinct clades corresponding to taxonomic families, although not fully mirroring the distribution in the general species phylogenetic tree. ACW-1 homologues in Glomerellaceae and Xylariaceae were grouped under the same clade, just like in the general species tree, but Nectriaceae sequences fell into a different clade. Aspergillaceae sequences were placed in a more distant clade. Saccharomycetaceae ACW-1 paralogs were grouped into two distinct subclades. B) Maximum-likelihood phylogenetic reconstruction of NCW-3 homologues displayed a similar pattern. They were distributed into thirteen clades consistent with family-level taxonomy. However, homologues from Podosporaceae, Chaetomiaceae, Lasiosphaeriaceae and Sordariaceae were segregated more distantly than in the species tree. Paralogs frequently occupied distinct subclades and did not group together. Sequences were aligned with MAFFT [112], edited in Jalview v2 [114] and reconstructed into maximum-likelihood (ML) phylogenetic trees using the PROTGAMMAIWAG model in RAxML [115]. Phylogenetic trees were visualized and annotated in iTOL v7.2.2 [116].

NCW-3 homologues showed a similar pattern. They formed 13 clades largely consistent with family-level taxonomy (**Figure 2B**), although some (*Podosporaceae*, *Chaetomiaceae*, *Lasiosphaeriaceae* and *Sordariaceae*) were distributed differently than in the species-level phylogeny (**Figure 1B**). In the latter, *Sordariaceae*, *Chaetomiaceae*, *Podosporaceae* and *Lasiosphaeriacea* were closely related and part of the same subclade. In contrast, the NCW-3 homologues in *Podosporaceae* and *Chaetomiaceae* were distributed farther from those of *Sordariaceae* and *Lasiosphaeriaceae*. Unlike ACW-1, NCW-3 paralogs failed to group together.

To quantitatively assess the evolutionary patterns suggested by these comparative analyses, Shannon entropy was calculated to assess sequence conservation across both protein families. Median Shannon entropy values were 0.39 (IQR: 0.25-0.60) for ACW-1 homologues and 0.40 (IQR: 0.25-0.64) for NCW-3 homologues, indicating moderate conservation and no significant difference between groups (Mann-Whitney U test, p-value > 0.05). To further evaluate relative evolutionary constraints, non-synonymous-to-synonymous replacement rate ratios (dN/dS) were estimated using the M0 model of codon substitution implemented in codeml from the Phylogenetic Analysis by Maximum Likelihood (PAML) package [41]. To minimize bias associated with the broader phylogenetic distribution of ACW-1, this comparison was restricted to shared taxa within Sordariales, where *N. crassa* is classified. Both proteins showed similar dN/dS ratios of less than 1 (ω= 0.1495 for ACW-1 and ω= 0.1482 for NCW-3), which is consistent with purifying selection acting on both protein families, suggesting removal of deleterious mutations and preservation of protein function. Median Shannon entropy values calculated within Sordariales were 0.46 (IQR: 0.27-0.68) for ACW-1 and 0.42 (IQR: 0.24-0.65) for NCW-3 homologues, further supporting similar (Mann-Whitney U test, p-value > 0.05) moderate conservation within this lineage. Together, these results suggested that ACW-1 and NCW-3 family evolution is decoupled from species-level phylogenetic relationships. However, both proteins shared similar global evolutionary constraints and, within Sordariales, also displayed similar signatures of purifying selection and moderate conservation, consistent with the presence of functional constraints.

### ACW-1 homologues exhibited higher sequence divergence in Saccharomycotina yeasts

ACW-1 and NCW-3 homologues were also clustered through a sequence similarity network (SSN) [42] to identify potential functional or structural similarities, as well as to independently assess the degree of sequence divergence within and across taxonomic lineages (**Figure 3, S1** and **S2**). ACW-1 homologues were analyzed across identity thresholds ranging from 60 to 99% (data not shown). At low thresholds (60-65%), homologues collapsed into broad groups that obscured lineage-level structure. Thresholds between 70-75% resolved major fungal lineages (*Daldinia*, *Mortierellaceae*, Serinales, Saccharomycotina). At high thresholds (above 90%), excessive fragmentation occurred, separating homologues from the same taxonomic orders into multiple subgroups. A threshold of 85% was selected as it split homologues into 9 biologically consistent clusters and 14 singletons (which were discarded for subsequent analyses) without overfragmentation (**Figure 3A**). At this threshold, cluster 1 included the majority of ACW-1 homologues (82%), including representatives from multiple filamentous fungi lineages. In contrast, Saccharomycotina sequences were distributed across distinct clusters (clusters 2 and 3), each containing paralogues from multiple yeast families. Smaller clusters corresponded to *Lipomycetaceae* (cluster 4), Serinales (clusters 5 and 8), *Mortierellaceae* (Mucuromycota) (cluster 6) and Dipodascales (cluster 7). This clustering pattern was consistent with the ACW-1-based phylogeny (**Figure 2A**), in which Saccharomycotina sequences formed the most separated and dispersed clades. Together, these results indicated that ACW-1 homologues display greater sequence divergence within Saccharomycotina yeasts compared to filamentous fungi, where homologues largely remained within a single major sequence cluster (cluster 1, **Figure 3A**).

**Figure 3.**
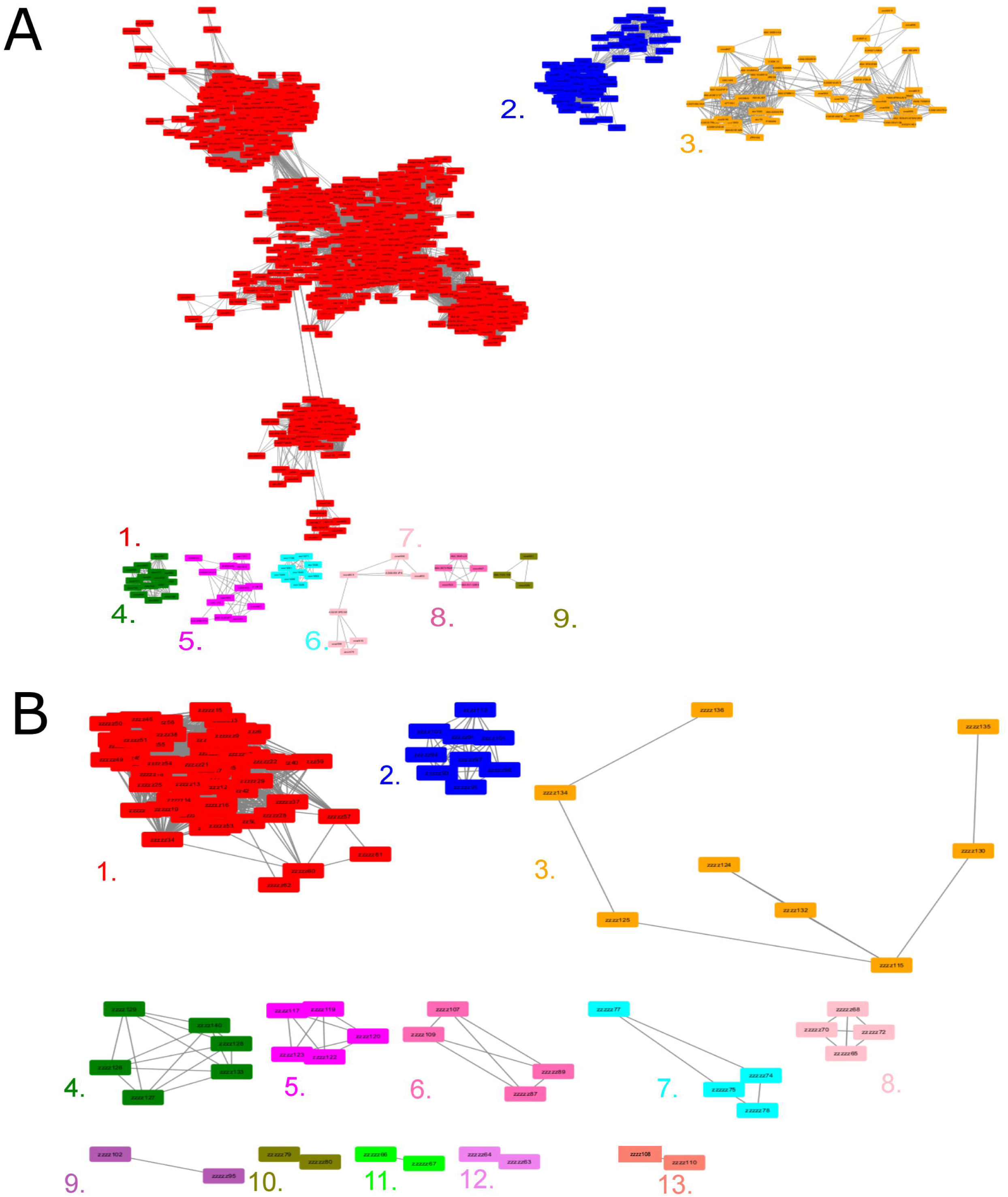
ACW-1 diversification occurred mainly in yeasts, while NCW-3 diversified more homogeneously. A) Protein clustering analysis of ACW-1 homologues at an 85% sequence identity threshold yielded nine distinct clusters. Most ACW-1 homologues (82%), mainly found in filamentous fungi, grouped in Cluster 1. The remaining clusters comprised mostly yeast sequences. Clusters 2 and 3 featured Saccharomycotina paralogs. Cluster 4 included Lipomycetaceae homologues. Clusters 5 and 8 grouped Serinales homologues. Cluster 7 featured Dipodascales homologues. Cluster 6 contained homologues from Mortierellaceae, which belongs to another subphylum (Mucoromycotina). B) Protein clustering analysis of NCW-3 homologues at a 75% sequence identity threshold produced thirteen clusters. Unlike ACW-1, NCW-3 homologues did not cluster around a single group, but were distributed more evenly across clusters defined largely by taxonomic families, including Sordariomycetidae (clusters 1 and 3), Saccotheciaceae (cluster 2), Sordariales (cluster 4), Phyllostictaceae (cluster 5), Botryosphaeriaceae (cluster 6), Cordycipitaceae (cluster 7), Pyriculariaceae (cluster 8). Trypetheliaceae (cluster 9), Mycosphaerellaceae (cluster 10), Schizotheciaceae (cluster 11), Ophiostomataceae (cluster 12), and Venturiaceae (cluster 13). Protein clustering analyses were performed in EFI-EST [42] and visualized in Cytoscape v3.10.3 [111].

To explore whether structural features were conserved despite sequence divergence, a representative mature ACW-1 homologue from each SSN cluster was modeled in AlphaFold2 [43] (**Figure S1A**). All models shared a conserved core architecture characterized by a β-sheet structure and a predicted C-terminal intrinsically disordered region (IDR, in magenta). The predicted structure of ACW-1 is addressed more comprehensively later on. Consensus sequence analysis further revealed conserved motifs across all clusters (**Figure S1B**). Pairs of cysteine residues (pink, arrow) were conserved near the N- and C-termini, consistent with potential disulfide bonds formed between the predicted α-helices and their closest β-sheet. Asparagine residues arranged in xNNN motifs near the middle of the sequences and NxS/T motifs were also present in all clusters (green, red dotted boxes), the latter suggesting the preservation of N-glycosylation sites potentially needed for structure or function. A conserved hydrophobic core enriched in leucine residues was also detected across the lengths of the proteins (blue, arrows; mapped to **Figure S1A** in yellow). These features suggested conservation of a shared structural scaffold despite pronounced sequence divergence among ACW-1 homologues.

### NCW-3 diversified more homogeneously than ACW-1

NCW-3 homologues were analyzed across identity thresholds ranging from 30 to 99% (data not shown). At lower thresholds (30-50%), homologues clustered in a small number of broad groups which masked taxonomic structure. Thresholds between 65-70% resolved major fungal lineages, including Sordariomycetidae, Dothideomycetes and Agaricales. At higher thresholds (above 80%), sequences were split into family-level groups but with a substantial proportion of singletons (24% of the sequences), which, given the small number of homologous sequences analyzed (139), overfragmented data and reduced interpretability. A threshold of 75% identity was selected as a balance between resolution and fragmentation, yielding 13 taxonomically consistent clusters and 18% singletons (**Figure 3B**). At this threshold, most clusters corresponded to single taxonomic families or closely related lineages: Sordariomycetidae (clusters 1 and 3), *Saccotheciaceae* (cluster 2), Sordariales (cluster 4), *Phyllostictaceae* (cluster 5), *Botryosphaeriaceae* (cluster 6), *Cordycipitaceae* (cluster 7), *Pyriculariaceae* (cluster 8), *Trypetheliaceae* (cluster 9), *Mycosphaerellaceae* (cluster 10), *Schizotheciaceae* (cluster 11), *Ophiostomataceae* (cluster 12), and *Venturiaceae* (cluster 13). This clustering pattern was consistent with the NCW-3-based phylogeny (**Figure 2B**), where homologues largely grouped according to taxonomic family. Compared to ACW-1, NCW-3 homologues exhibited a more even distribution of sequence similarity across fungal lineages. This pattern suggests a more homogeneous distribution of sequence divergence across taxa, in contrast to the more lineage-expanded pattern observed for ACW-1 in Saccharomycotina yeasts.

To identify structural conservation patterns, representative mature NCW-3 homologues from each SSN cluster were modelled using AlphaFold2 [43] (**Figure S2A**). All models showed a highly conserved structure, characterized by a N-terminal β-hairpin and a C-terminal pore-like structure. Consensus sequences analysis across clusters (**Figure S2B**) revealed strong conservation of the N-terminal region corresponding to the predicted β-hairpin, including motifs such as CG, LCPx, ACYS and YxC, previously described in the CBM-52 domain of the *S. pombe* Eng1 [36]. This CBM-52 domain has affinity for β-1,3-glucan and is necessary to localize Eng1 to the septa. Four conserved cysteine residues located in the middle and C-terminal regions (pink, arrows) were identified across all clusters and predicted to participate in disulfide bond formation to stabilize the β-hairpin structure. A conserved VPGGQ motif (red dotted box) was identified across all clusters too. These conserved structural and sequence features indicated strong structural constraints acting on NCW-3 homologues across fungal lineages, despite variation in sequence similarity among taxonomic family groups.

Together, these results suggested distinct evolutionary dynamics for ACW-1 and NCW-3. ACW-1 homologues displayed greater sequence divergence across broad fungal lineages, especially among Saccharomycotina yeasts. In contrast, NCW-3 homologues showed lower overall sequence divergence than ACW-1 but exhibited stronger clustering by taxonomic family and less lineage-specific expansion. Both proteins retained conserved predicted structural frameworks across clusters, supporting homology within the gene families.

### ACW-1 is built of leucine-rich repeats (LRR) modules that evolved at different rates

To investigate the structural organization of ACW-1 homologues, *N. crassa* (Nc) ACW-1 was used as a representative model. The predicted mature structure consisted of an elongated arrangement of right-handed β-sheets-rich elements forming concave and convex surfaces (**Figure 4A**). This core was capped by N- and C-terminal α-helices, which contained the highly conserved cysteine pairs predicted to stabilize the structure through disulfide bonds (**Figure S1B**). This structure was consistent with the predicted structures of ACW-1 homologues (**Figure S1A**). A predicted N-terminal Ecm33 domain (PF12454) (**Figure 4A**, orange) and a C-terminal disordered region (**Figure 4A**, magenta) were identified. Intrinsic disorder was confirmed with ESpritz [44], AIUPred [45], IUPred2A [46], identifying a C-terminal intrinsically disordered region (IDR) of 54 amino acids length (∼200 Å, calculated as the distance along alpha carbon atoms in PyMOL v3 [47] (**Figure 4B**). The elongated structure featured a hydrophobic core (**Figure 4C**, yellow) interspersed with polar (green), positively charged (pink) and negatively charged (blue) amino acids. This pattern was consistent with the conservation of hydrophobic amino acids, particularly leucine, across ACW-1 homologues (**Figure S1B**), suggesting the presence of leucine-rich repeat (LRRs) regions. LRRs are formed by 20-30 amino acids that contain the hallmark motif (LxxLxLxxN/CxL), where x can be any amino acid and leucine positions can be occupied by valine, isoleucine and phenylalanine, whereas asparagine positions can be substituted by threonine, serine or cysteine residues [48,49]. The repeat also includes segments of variable composition and length. Structurally, LRRs-containing proteins adopt a horseshoe shape, where the LRR motif makes up the concave face as a series of β-sheets and their adjacent loops, while the length-variable segments are located on the convex face and form other secondary structures [50]. LRRs-containing proteins are widespread in prokaryotes and eukaryotes. They are involved in protein-protein interactions [50], including signal transduction [51], cell adhesion [52], innate immunity [53] and gene regulation [54], among others.

**Figure 4.**
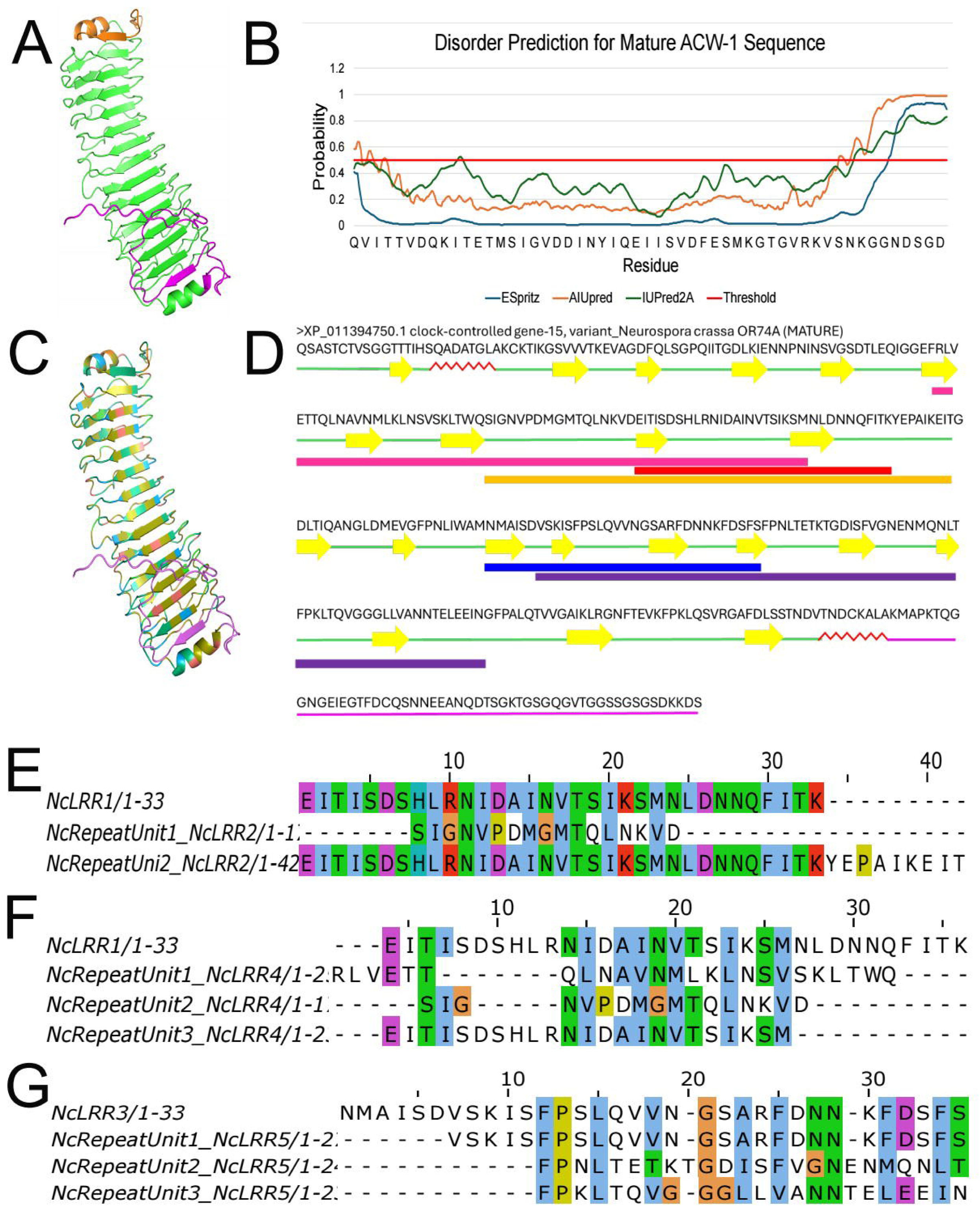
ACW-1 is built of leucine-rich repeats (LRR) that evolved at different rates. A) The mature ACW-1 structure was predicted as an elongated β-helix flanked by N- and C-terminal α-helices. An N-terminal Ecm33 domain (orange) and a C-terminal disordered region were also inferred (magenta). B) C-terminal intrinsic disorder was confirmed by ESpritz [44], AIUpred [45] and IUPred2A [46]. C) ACW-1 displayed a predominantly hydrophobic core (yellow) interspersed with polar (green), positively charged (pink) and negatively charged (blue) residues. D) Five significant LRR hits (c-E-value < 0.05) of varying lengths were identified across ACW-1, several of which overlapped: Nc LRR1 (residues 122-154, red), Nc LRR2 (residues 105-163, orange), Nc LRR3 (residues 187-219, blue), Nc LRR4 (residues 80-144; pink) and Nc LRR5 (residues 193-266, purple), listed in descending order of confidence prediction. All formed a concave structure made of hydrophobic β-sheets with adjacent loops, while the structural configuration of the convex surface varied. E) Examination of Nc LRR2 and F) Nc LRR4 consisted of two and three divergent repeats of Nc LRR1, respectively, whereas G) Nc LRR5 comprised three divergent repeats of Nc LRR3. Structural predictions were generated with AlphaFold2 [43] and visualized in PyMOL v3 [47]. Nc LRRs were aligned with MAFFT [112] through the Galaxy server [113] and visualized in Jalview v2 [114].

Because tandem repeats are common in fungal CWPs, ACW-1 and its homologues were screened for LRR domains using hmmscan [55]. As expected for highly divergent repeat sequences like LRRs [56], multiple LRR domains were detected with low but consistent statistical support (**Table S3**), showing independent E-values (i-Evalues) between 3.2x10^-6^ and 1x10^3^, and conditional E-values (c-Evalues) between 3.4x10^-9^ and 1x10^1^, including 13 predicted LRR domains in ACW-1. Since c-Evalues measure the statistical significance of a specific domain once true homology has been established [57], and structural conservation supported homology among ACW-1 homologues (**Figure S1A**), c-Evalues below 0.05 were used as the threshold for LRR domain significance. Five statistically supported LRR domains of varying lengths were predicted across the mature sequence of ACW-1 (in order of statistical support, **Figure 4D**): Nc LRR1 (residues 122-154, red), Nc LRR2 (residues 105-163, orange), Nc LRR3 (residues 187-219, blue), Nc LRR4 (residues 80-144, pink), and Nc LRR5 (residues 193-266, purple). Long predicted annotations (Nc LRR2, Nc LRR4, NcLRR5) were hypothesized to represent composite regions formed by the duplication and fusion of shorter regions (Nc LRR1, Nc LRR3). To resolve this, self-alignment and predicted overlapping LRR domain boundary comparisons were used to define operational putative LRR units that could serve as evolutionary units for further analyses (**Figure 4E-G**). Nc LRR2 was hypothesized to be composed of 2 divergent Nc LRR1 units **(Figure 4E**), where the first one (NcRepeatUnit1_NcLRR2) was highly divergent and degenerate, and the second (NcRepeatUnit2_NcLRR2) was similar to Nc LRR1 but with a few additional residues. Nc LRR4 overlapped with Nc LRR2 and was also hypothesized to be composed of 3 divergent Nc LRR1 units (**Figure 4F**), with the first one (NcRepeatUnit1_NcLRR4) being slightly divergent and shorter, the second one (NcRepeatUnit2_NcLRR4) was a shorter, degenerate sequence shared by Nc LRR2, and the third (NcRepeatUnit3_NcLRR4) was shorter version of Nc LRR1. Nc LRR5 was composed of one shorter Nc LRR5 unit (NcRepeatUnit1_NcLRR5) and 2 variations (**Figure 4G**) which were shorter, slightly divergent versions of Nc LRR3 (NcRepeatUnit2_NcLRR5 and NcRepeatUnit3_NcLRR5). These results suggested that overall 6 putative LRR units (ACW1_RU1, ACW1_RU2, ACW1_RU3, ACW1_RU4, ACW1_RU5, ACW1_RU6) were present in ACW-1, organized into 2 larger LRR regions corresponding to Nc LRR4 and Nc LRR5 (**Figure 5A**).

**Figure 5.**
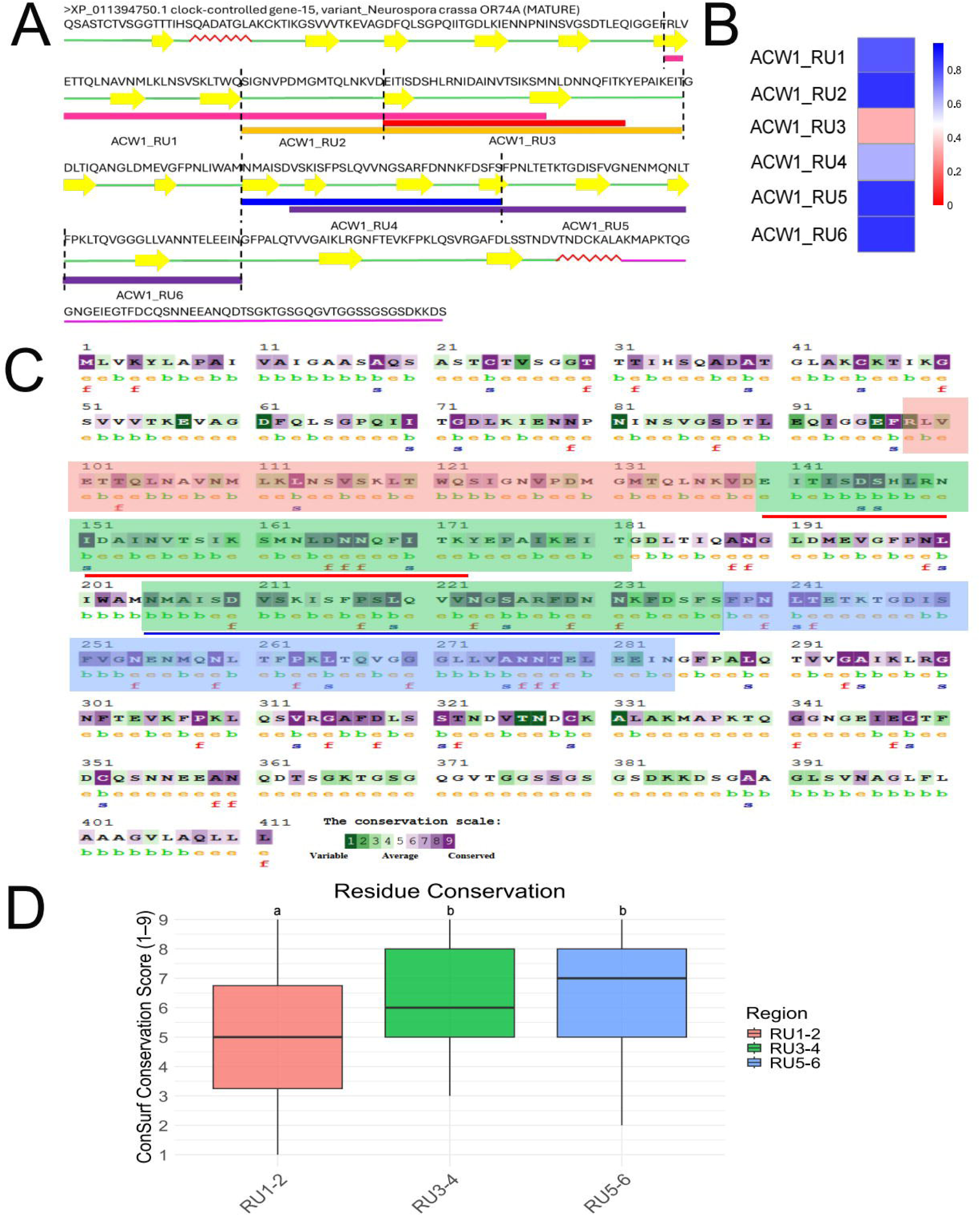
ACW-1 showed asymmetric evolutionary pressures along its sequence. A) Predicted long LRR domains, Nc LRR2, Nc LRR4 and NcLRR5, were composed of six LRR repetitive units (RU): ACW1_RU1, ACW1_RU2, ACW1_RU3, ACW1_RU4, ACW1_RU5, and ACW1_RU6. B) Pairwise distances (p-distances) between the ancestral reconstructed sequence and these six LRR RUs were calculated and revealed a gradient of increasing divergence toward the protein termini. Higher values (blue) indicate greater divergence, and lower values (red) indicate higher similarity. C) Amino acid conservation analysis revealed distinct conservation patterns. Annotations “s” and “f” indicate predicted structural and functional residues, respectively. Conserved residues are shown in purple and variable positions appear in green. Nc LRR1 (red underline) and Nc LRR3 (blue underline), are shown for context. Three regions were determined positionally for conservation analysis: region RU1-2 was close to the N-terminus and composed of ACW1_RU1 and ACW1_RU2 (salmon pink shadow); region RU2-4 localized in the middle of ACW-1 and included ACW1_RU3 and ACW1_RU4 (green shadow), and region RU5-6 was close to the C-terminus and composed of ACW1_RU5 and ACW1_RU6 (blue shadow). D) Residue conservation varied among these three regions; regions RU3-4 and RU5-6 showed higher median conservation scores than region RU1-2 (Kruskal Wallis test, P-value < 0.05). Statistical analyses were performed in RStudio v4.4.2 [110]. The scored conservation and predicted functional or structural roles analysis was performed with ConSurf [61].

To estimate the degree of divergence among these 6 putative LRR units (RUs), an ancestral LRR sequence was reconstructed. Statistically significant predicted LRR hits (c-Evalue < 0.05) considerably varied in length across ACW-1 and its homologues, with some annotations extending up to 266 amino acids. While canonical LRR motifs typically span 20-30 residues, within the context of ACW-1 and its homologues most predicted LRR segments were approximately 35 amino acids in length (**Figure S3A**). Based on this observation, predicted LRR domains ranging from 20 to 40 amino acids were extracted and used for ancestral sequence reconstruction using IQ-TREE v3 (**Figure S3B**) [58–60]. This set included Nc LRR1 (red) and Nc LRR3 (blue). The reconstructed ancestral sequence was inferred as EVTISDTKLRSIDAINVTSISSMNLDNNQFITKFSLSS.

Pairwise distances (p-distances) were calculated between this reconstructed ancestral sequence and the modern ACW-1 putative LRR units (**Figure 5B**), where lower values indicated greater similarity and higher values indicated increased divergence. This analysis revealed a divergence gradient along ACW-1. The putative repeat unit located near the middle of the protein, ACW1_RU3, which corresponded largely to Nc LRR1 with a few more amino acids, exhibited the lowest divergence from the ancestral sequence (p-distance = 0.33). Consistently, Nc LRR1 was also a direct descendant of the ancestral sequence in the reconstructed ancestral phylogeny (**Figure S3B**, red; branch length = 0.17). In contrast, divergence increased toward both termini. Toward the N-terminus, ACW1_RU2 and ACW1_RU1 showed p-distances of 0.86 and 0.79, respectively. Toward the C-terminus, ACW1_RU4, ACW1_RU5 and ACW1_RU6 displayed p-distances of 0.62, 0.86 and 0.86, respectively. These results suggested a gradient of sequence divergence from the central region of ACW-1 towards its N- and C-termini.

To assess conservation patterns among modern ACW-1 homologues independently of the ancestral reconstruction, an amino acid scored conservation analysis as well as prediction of structural and functional roles was performed using ConSurf [61]. For this purpose, ACW1_RUs were grouped into three regions based on their position within ACW-1. The N-terminal region (RU1-2) comprised ACW1_RU1 and ACW1_RU2 (**Figure 5C**, red), the central region (RU3-4) contained ACW1_RU3 and ACW1_RU4 (**Figure 5C**, green), and the C-terminal region RU5-6 included ACW1_RU5 and ACW1_RU6 (**Figure 5C**, blue). Statistical comparison of scores revealed distinct conservation patterns among these regions (**Figure 5D**). The N-terminal RU1-2 region exhibited a median conservation score of 5 (IQR: 3.25-6.75), corresponding to average conservation according to the ConSurf scale. In contrast, the central RU3-4 region (median score 6, IQR: 5-8) and C-terminal region RU5-6 (median score 7, IQR: 5-8) showed statistically higher median scores, indicating stronger sequence constraint across extant homologues. Residues predicted by ConSurf to be functionally important (annotation “f”, **Figure 5C**) were predominantly among the most conserved positions (scores 8-9).

Experimental studies have identified 2 N-glycosylation sites in ACW-1, N155 (for context, located on ACW1_RU3) and N240 (located on ACW1_RU4) [28]. These glycosylations are processed by the α-1,6-mannanases DFG5 and DCW1 [14] and subsequently cross-linked to β-1,3/1,4 mannans [14,28,29,34] by transglycosidases GEL-1, GEL-2, and GEL-5 [35] to integrate ACW-1 into the cell wall. Predictions using NetNGlyc v1.0 [62] identified additional potential N-glycosylation sites toward the C-terminus (N223, N240, N259, N276, N301). These asparagine residues were also predicted by ConSurf to be highly conserved (above 7) and frequently annotated as function-associated (annotated as “f”).

Despite measuring different evolutionary properties, ancestral p-distance estimates and ConSurf conservation scores revealed a parallel convergent pattern along ACW-1, in which the central region (RU3-4) displayed both reduced divergence from the reconstructed ancestral sequence (as exemplified by ACW1_RU3) and also increased conservation across extant homologues. In both approaches, the N- terminal region was the most divergent, whereas the C-terminal region showed both the greatest historical divergence and highest conservation in extant homologues, consistent with differential evolutionary constraints across ACW-1 regions. Together, these results suggest asymmetric evolutionary pressures along ACW-1, in which distinct regions have followed different evolutionary trajectories shaped by both historical divergence and present-day functional requirements. A comparable regional divergence pattern was not detected in NCW-3 under the same analytical framework (data not shown).

### Deletion of NCW-3 showed more pronounced morphological and growth defects than ACW-1 in *N. crassa*

Although ACW-1 and NCW-3 are both predicted to associate with cell wall β-1,3-glucans via different mechanisms [34,36], their functional roles were evaluated using homokaryotic *N. crassa* knock-out (KO) mutants. The KO mutants for the genes of each protein, Nc Δ*acw-1* and Nc Δ*ncw-3*, and a wild-type (WT) strain were analyzed for morphology, growth rate, branching pattern and conidiation (**Figure 6**). KO mutants were molecularly confirmed to verify their homokaryotic status and relate any observed effect to the absence of the respective CWP gene (**Supplementary Material and Figure S4**). Under standard growth conditions, the KO mutants showed a visually distinct macroscopic gross colony architecture in contrast with the WT strain (**Figure 6A-C**). The Nc WT strain colony resembled an even mycelial mat with well-defined aerial hyphae radiating out and surrounding a central pigmented conidial mass (**Figure 6A**). In contrast, the Nc Δ*acw-1* strain colony appeared less uniform and more irregular in the aerial hyphae pattern, the central conidial mass looked reduced and less pigmented (**Figure 6B**). The Nc Δ*ncw-3* strain colony exhibited a smoother surface, less dense aerial hyphae and visible but less pigmented conidia (**Figure 6C**). Microscopically, there were no apparent differences between the hyphal morphology of the Nc WT strain (**Figure 6D**) and the Nc Δ*acw-1* strain (**Figure 6E**), and both strains exhibited long and cylindrical hyphae and dichotomous branching. Both the Nc Δ*acw-1* (median diameter: 2.73 μm, IQR: 2.99-2.27 μm) and Nc Δ*ncw-3* (median diameter: 2.28 μm, IQR: 2.81-1.87 μm) hyphae showed reduced diameters in contrast with the Nc WT strain (median diameter: 3.20 μm, IQR: 3.44-2.93 μm). Moreover, the Nc Δ*ncw-3* strain showed an aberrant phenotype characterized by shorter branches (median branch length: 77.2 μm, IQR: 113-45.7 μm) than Nc WT (median branch length: 115 μm, IQR: 179-53 μm) and Nc Δ*acw-1* (median branch length: 97.6 μm, IQR: 164-41.4 μm) (**Figure 6F**). There was not a significant difference between the vegetative growth (**Figure 6G**) of the Nc WT (2.10 ± 0.27 mm/h) and Nc Δ*acw-1* (2.38 ± 0.23 mm/h) strains, whereas it decreased in the Nc Δ*ncw-3* (1.50 ± 0.18 mm/h). No significant differences were observed in the hyphal ramification (**Figure 6H**) between Nc WT (14.3 ± 2.90 branches) and Nc Δ*acw-1* (13.6 ±3.78 branches), while it decreased in Nc Δ*ncw-3* (11.5 ± 3.54 branches) strain. Conidiation decreased in the KO mutant strains in comparison with the Nc WT strain (**Figure 6I**). The Nc Δ*acw-1* (4.60 ± 0.40 x 10^7^ conidia/mL) and the Nc Δ*ncw-3* (2.71 ± 0.60 x 10^7^ conidia/mL) strains produced less conidia than the Nc WT (1.10 ± 0.12 x 10^8^ conidia/mL) strain. These results suggested that ACW-1 and NCW-3 are not strictly required for normal morphology of the hyphae, growth rate and ramification. However, the altered macroscopic gross colony phenotype and reduced conidiation observed in both KO mutants pointed to defects in cell wall organization or remodeling, with the Nc Δ*ncw-3* strain exhibiting more pronounced phenotypic alterations than the Nc Δ*acw-1* strain.

**Figure 6.**
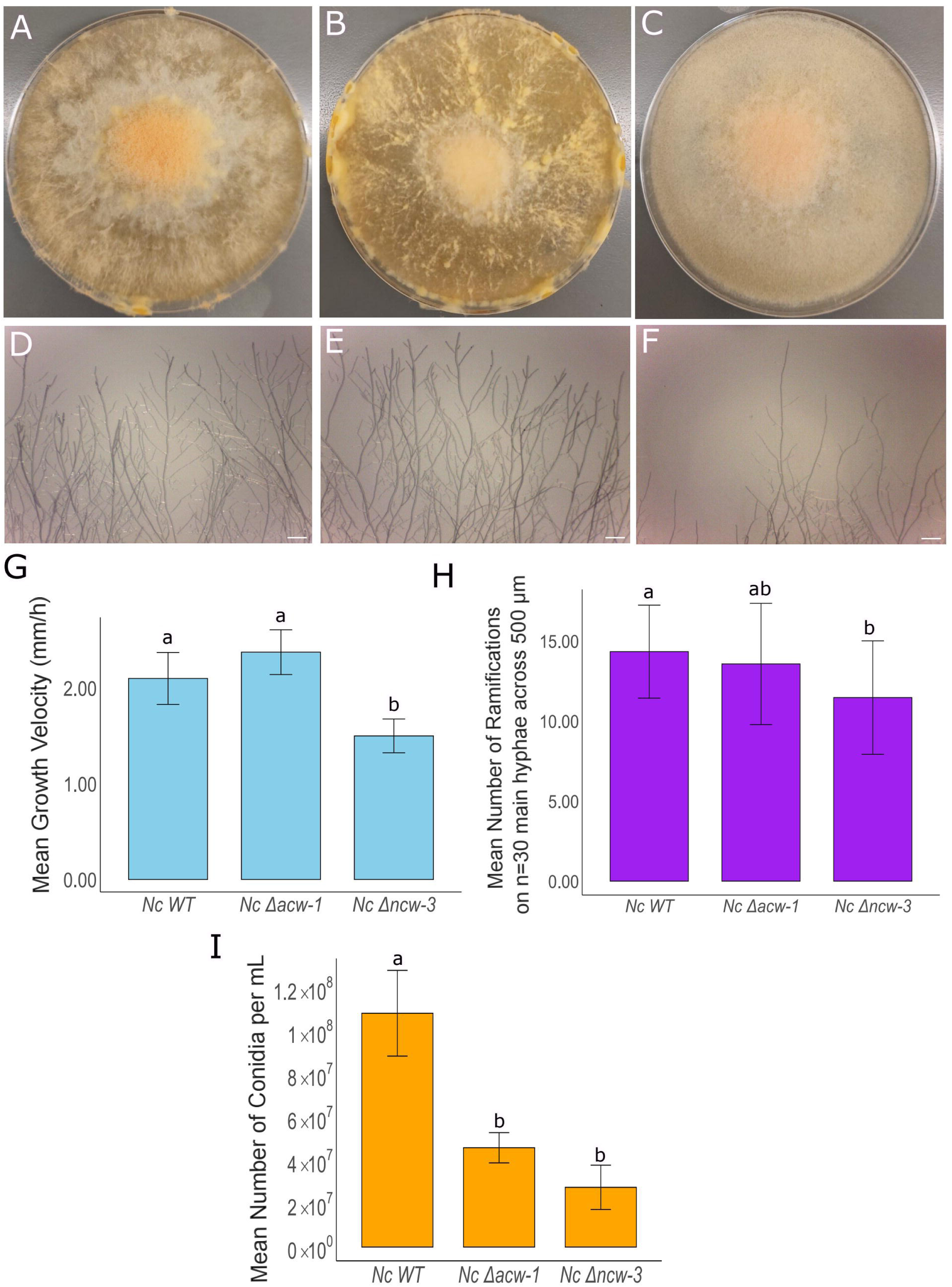
*N. crassa* Δ*ncw-3* showed a more pronounced morphological defect than *N. crassa* Δ*acw-1*, which remained largely similar to *N. crassa* WT. Morphological characterization of (A, D) *N. crassa* WT, (B, E) *N. crassa* Δ*acw-1*, and (C, F) *N. crassa* Δ*ncw-3* showed no differences between the first two, while *N. crassa* Δ*ncw-3* showed reduced hyphal density and diameter, and shorter branches. Bright field images shown are representatives of 10 fields; they were obtained in Leica EZ4 stereomicroscope (Leica Microsystems) and edited in Fiji [127]. Scale bar, 40 μm. G) Vegetative growth rate and (H) branching of the three strains were quantified showing no differences between Nc WT and Nc Δ*acw-1*, while Nc Δ*ncw-3* showed a statistically significant reduction of growth rate and branching. I) Conidiation decreased in Nc Δ*acw-1* and Nc Δ*ncw-3* strains. Growth rate measurements were the mean of 5 replicates. Branching frequency was quantified as the mean of 30 main hyphae over a length of 500 μm. Conidiation measurement was the mean of 3 replicates. Mean parameters are shown, error bars represent standard deviation. Comparison between strains was performed by one-way ANOVA. Statistical significance was determined at P-value < 0.05. The same letter indicates a non-significant statistical difference, while different letters indicate a significant statistical difference. Statistical analyses were conducted in RStudio v4.4.2 [110] (RStudio Team, 2020).

### The *N. crassa* **Δ***acw-1* strain exhibited the highest susceptibility to CFW, a chitin synthesis inhibitor

To assess the impact of ACW-1 and NCW-3 loss on the *N. crassa* cell wall, the Nc Δ*acw-1* and Δ*ncw-3* strains were subjected to stress susceptibility assays with Calcofluor White (CFW) and Congo Red (CR), dyes that interfere with the synthesis of cell wall chitin and β-1,3-glucan, respectively [63–65] (**Figure 7**).

**Figure 7.**
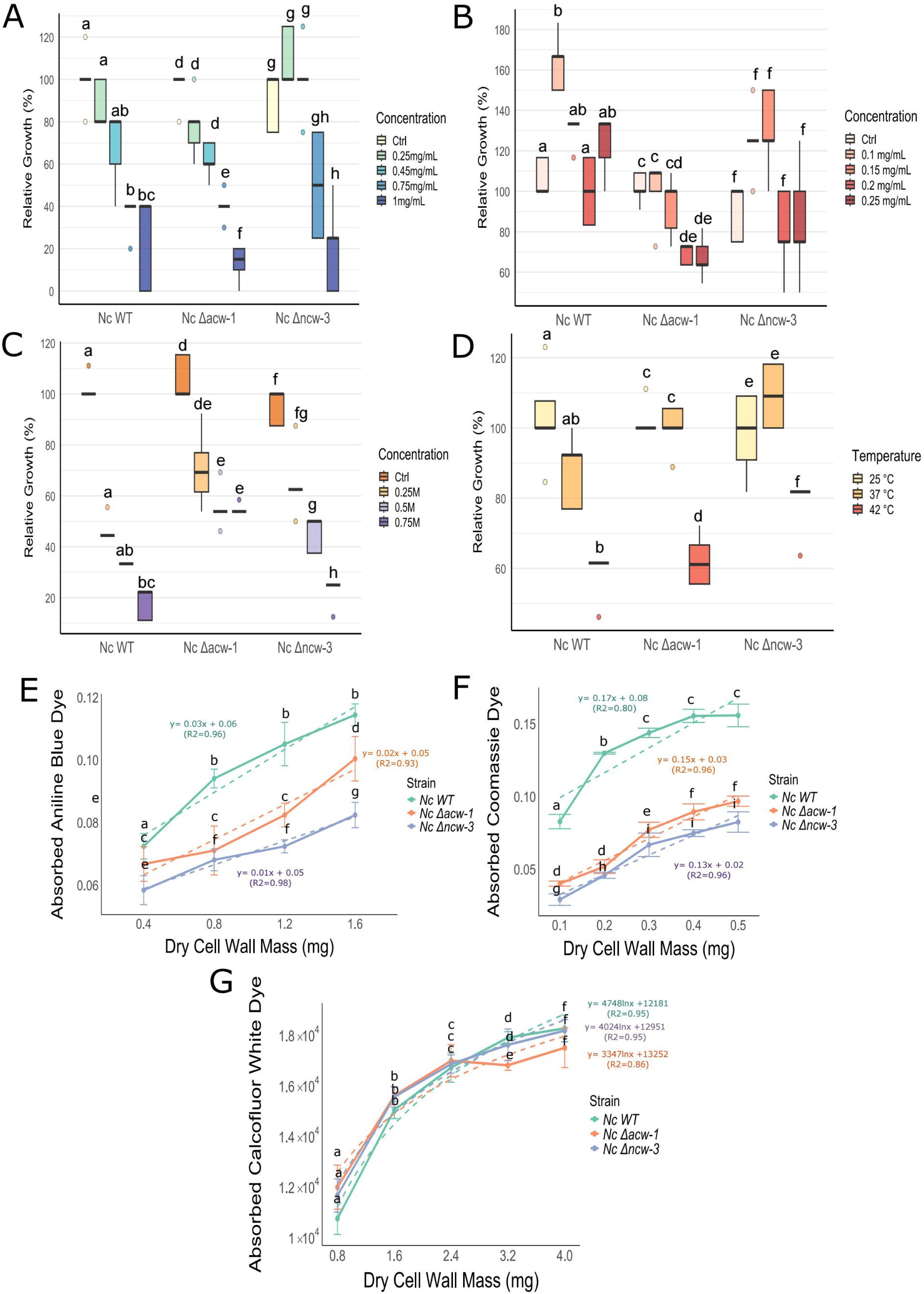
ACW-1 and NCW-3 play similar roles in stabilizing cell wall. β-1,3-glucans and are involved in the maintenance of the integrity of the *Neurospora crassa* cell wall. Nc WT, Nc Δ*acw-1* and Nc Δ*ncw-3* strains were grown in A) 0.25-1 mg.mL^-1^ CFW, B) 0.1-0.25 mg.mL^-1^ CR, C) 0.25-0.75 M NaCl, and D) 37 °C and 42 °C to assess any cell wall defect affecting its stability. Nc Δ*acw-1* and Nc Δ*ncw-3* strains displayed increased sensitivity to CFW and CR but more tolerance to osmotic stress than the Nc WT strain. The Nc Δ*ncw-3* strain also showed increased temperature tolerance at 42 °C. Data represent the median relative growth (%) from 5 replicates per condition. Error bars show interquartile range (IQR). Kruskal-Wallis tests with post *hoc* Dunn tests (P-value < 0.05) were used to compare growth between control and stressor levels within each strain. Matching letters denote non-significant statistical differences; different letters indicate significant statistical differences. E) Aniline blue, F) Coomassie Blue and E) CFW dye binding assays were used to quantify β-1,3-glucans, total cell wall protein content and chitin, respectively, in purified cell walls of the Nc WT, Nc Δ*acw-1* and Nc Δ*ncw-3* strains. The Nc Δ*acw-1* and Nc Δ*ncw-3* strains showed decreased levels of β-1,3-glucans and total cell wall protein compared with the Nc WT strain, while chitin contents remained unchanged. Mean absorbed dye values from 3 replicates are shown, calculated as the difference between fluorescence intensity measurements in the supernatant, representing unbound dye, and controls lacking cell wall material. Error bars represent standard deviation. One-way ANOVAs (P-value < 0.05) were conducted to compare differences of absorbed dye across dry cell wall amounts. Mathematical model equations that describe the relationship between absorbed dye and dry cell wall mass are shown. Linear models best described aniline and Coomassie dyes absorption, whereas CFW absorption followed a logarithmic model. Statistical analyses were performed in RStudio v4.4.2 [110].

In the CFW inhibition assay (**Figure 7A**), WT median growth decreased from 100% (IQR: 100-100%) in the control condition without CFW (Ctrl) to 80% (IQR: 80-100%) at 0.25 mg/mL CFW and remained stable at 80% (IQR: 60-80%) at 0.45 mg/mL CFW. Growth was further reduced at higher CFW concentrations, reaching 40% (IQR: 40-40%) at 0.75 mg/mL CFW and remaining at 40% (IQR: 0-40%) at 1.00 mg/mL CFW. The median growth of the Nc Δ*acw-1* strain decreased to 80% (IQR: 70-80%) at 0.25 mg/mL CFW. It dropped to 60% (IQR: 60-70%) at 0.45 mg/mL CFW. At higher CFW concentrations, it declined to 40% (IQR: 40-40%) at 0.75 mg/mL CFW, and to 15% (IQR: 10-20%) at 1.00 mg/mL CFW. In contrast, the Δ*ncw-3* strain showed no significant reduction in growth at 0.25 mg/mL CFW (100%, IQR: 100-125%) and 0.45 mg/mL CFW (100%, IQR:100-100%). It decreased to 50% (IQR: 25-75%) at 0.75 mg/mL CFW, and further declined to 25% (IQR: 0-25%) at 1.00 mg/mL CFW. The increased sensitivity of the Nc Δ*acw-1* and Nc Δ*ncw-3* strains to CFW pointed to defects or weakening in the cell wall, exacerbated by the addition of CFW. These results suggested that ACW-1 and NCW-3 may contribute to cell wall integrity, with ACW-1 playing a slightly more prominent role on this than NCW-3.

### The *N. crassa* **Δ***acw-1* strain exhibited the highest susceptibility to CR, a glucan synthesis inhibitor

In the CR inhibition assay (**Figure 7B**), Nc WT median growth increased to 167% (IQR: 150-167%) at 0.1 mg/mL CR, then decreased to 133% (IQR: 133-133%) at 0.15 mg/mL CR. At 0.20 mg/mL CR, median growth was statistically similar to the control condition without CR (100%, IQR: 83-117%). At 0.25 mg/mL CR, it increased again to 133% (IQR: 117-133%), though this increase was not statistically different to the control condition. The median growth of the Nc Δ*acw-1* strain remained statistically similar in the control (100%, IQR: 100-109%), 0.10 mg/mL (109%, IQR: 100-109%) and 0.15 mg/mL CR (100%, IQR: 82-100%) concentrations. At 0.20 mg/mL CR, median growth rate decreased to 73% (IQR: 67-73%) and further declined to 64% (IQR: 64-73%) at 0.25 mg/mL CR. The median growth rate of the Nc Δ*ncw-3* strain increased to 125% (IQR: 125-125%) at 0.10 mg/mL CR and remained statistically similar (125%, IQR: 125-150%) at 0.15 mg/mL CR. At 0.20 mg/mL, it decreased to 75% (IQR: 75-100%), and remained statistically similar (75%, IQR: 75-100%) at 0.25 mg/mL CR. Considering all tested inhibitory concentrations, both Nc Δ*acw-1* and Nc Δ*ncw-3* strains showed higher susceptibility to CR than Nc WT. In contrast with the Nc Δ*acw-1* strain, the growth inhibition of Nc Δ*ncw-3* showed a more plateaued response with increasing CR concentration and progressive β-1,3-glucan synthesis inhibition, which suggested a distinct functional contribution from that of ACW-1 to the integrity of the cell wall.

### The *N. crassa* Δ*acw-1* and Δ*ncw-3* strains exhibited increased resistance to NaCl-mediated osmotic stress

To assess the cell wall stability of Nc Δ*acw-1* and Nc Δ*ncw-3* strains under general stressors, NaCl-mediated osmotic stress and temperature inhibition assays were performed. In the NaCl-mediated osmotic stress inhibition assay (**Figure 7C**), Nc WT median growth rate decreased to 44% (IQR: 44-44%) at 0.25 M NaCl. It further decreased to 33% (IQR: 33-33%) at 0.50 M NaCl and declined to 22% (IQR: 11-22%) at 0.75 M NaCl. In contrast, the median growth rate of the Nc Δ*acw-1* strain decreased to 69% (IQR: 62-77%) at 0.25 M NaCl. It further decreased to 54% (IQR: 54-54%) at 0.50 M NaCl and remained stable (54%, IQR: 54-54%) at 0.75 M NaCl. The median growth rate of the Nc Δ*ncw-3* strain decreased to 62% (IQR: 62-63%) at 0.25 M NaCl. Growth dropped by 50% (IQR: 38-50%) at 0.50 M NaCl and declined to 25% (IQR: 25-25%) at 0.75 M NaCl. Both the Nc Δ*acw-1* and Nc Δ*ncw-3* strains exhibited increased resistance to NaCl-mediated osmotic stress, with the Nc Δ*acw-1* strain showing a more stable growth profile across increasing NaCl concentrations.

### The *N. crassa* Δ*ncw-3* strain exhibited tolerance to high growth temperatures

In the growth temperature assay (**Figure 7D**), Nc WT median growth rate decreased to 92% (IQR: 77-93%) at 37 °C, although the difference with growth at 25 °C (100%, IQR: 100-108%) was not statistically significant. However, the growth rate declined to 62% (IQR: 62-62%) at 42 °C. On the other hand, there was no significant difference between the Nc Δ*acw-1* strain growth at 25 °C (100%, IQR: 100-100%) and at 37 °C (100%, IQR: 100-106%). At 42 °C, growth declined to 61% (IQR: 56-67%). The median growth of the Nc Δ*ncw-3* strain increased to 109% (IQR: 100-119%) at 37 °C, although this difference was not statistically significant with growth at 25 °C (100%, IQR: 91-109%). Nevertheless, the growth rate declined to 82% (IQR: 82-82%) at 42 °C. The Nc Δ*ncw-3* strain displayed increased tolerance to elevated growth temperatures compared to the Nc WT and Nc Δ*acw-1* strains.

### Cell walls of *N. crassa* Δ*acw-1 and* Δ*ncw-3* strains showed reduced basal β-1,3 glucan and protein-associated signals, but unchanged basal chitin signal than the *N. crassa WT* strain

To evaluate the impact of ACW-1 and NCW-3 deletion on cell wall composition, dye-binding assays with purified cell walls were used as proxies for major cell wall components Aniline blue, which selectively binds β-1,3-glucan [66], was used to estimate relative β-1,3-glucan content. Coomassie brilliant blue, which interacts with proteins [67], was used to assess total cell wall protein content. Calcofluor White (CFW), a specific marker of fungal chitin [68], was used to evaluate chitin levels. The Nc WT strain showed an aniline blue signal of 0.03 Abs/mg dry cell wall mass. Both Nc KO mutants exhibited reduced binding with the Nc Δ*acw-1* strain showing 0.02 Abs/mg dry cell wall mass (66.67% of Nc WT aniline binding) and the Nc Δ*ncw-3* strain showing 0.01 Abs/mg dry cell wall mass (33.33% of Nc WT aniline binding) (**Figure 7E**).

A similar pattern was observed for total cell wall protein signal (**Figure 7F**). The Nc WT strain showed 0.17 Abs/mg dry cell wall. Both the Nc Δ*acw-1* strain (0.15 Abs/mg dry cell wall mass; 88.23% of Nc WT protein content), and the Nc Δ*ncw-3* strain (0.13 Abs/mg dry cell wall mass; 86.67% of Nc WT protein content) showed reduced binding.

In contrast, no substantial differences were observed in CFW binding across strains (**Figure 7G**). Together, these results indicated that deletion of ACW-1 and NCW-3 is associated with reduced β-1,3-glucan and protein signals in the cell wall of *N. crassa*, while chitin-associated signal remained unchanged. These changes are consistent with alterations in cell wall composition or organization in the KO strains.

### eGFP-V5-ACW-1 and NCW-3-V5-eGFP localized to distal septae and hyphal surface of *N. crassa*

Biochemical analyses suggest that both *N. crassa* ACW-1 and NCW-3 localize to the cell wall and interact with β-1,3-glucans [31], but their subcellular localization has not been directly characterized. To address this, ACW-1 and NCW-3 were N-(eGFP::V5::ACW-1) and C-terminal (NCW-3::V5::eGFP) tagged, respectively, with the enhanced green fluorescent protein (eGFP) and V5 epitopes and expressed in *N. crassa*. Expression assemblies for eGFP::V5::ACW-1 and NCW-3::V5::eGFP, *pMF272::NCACW-1* and *pMF272::NCNCW-3,* respectively, were designed and constructed (**Supplementary Materials,** and **Figure S5 and S6**). This configuration was aimed to prevent any secretion impairment since, in the case of ACW-1, eGFP::V5 tags were fused three amino acids downstream the N-terminal signal peptide cleaving site, far away from the C-terminal GPI addition site, and, in the case of NCW-3, eGFP::V5 tags were fused at the C-terminus, on the opposing side of the CBM-52 domain, which putatively interacts with β-1,3-glucans. Heterokaryotic *N. crassa* eGFP::V5::ACW-1 (Nc NCACW-1) and NCW-3::V5::eGFP (Nc NCNCW-3) expressing strains were generated, molecularly confirmed (**Supplementary Materials** and **Figure S6**) and analyzed in an inverted laser scanning confocal microscope (LSCM). Expression of eGFP::V5::ACW-1 and NCW-3::V5::eGFP fusions was also assessed by Western blot (**Supplementary Materials** and **Figure S7**). Nc NCACW-1 and Nc NCNCW-3 hyphae were analyzed at the active extension apical zone (within 30 μm from the tip) [69] and distal regions (beyond 120 μm from the tip) [70].

As expected, no fluorescence was detected in the distal (**Figure 8A**) or apical regions (**Figure 8B**) in the hyphae of the parental Nc FGSC9717 strain. In contrast, in the Nc NCACW-1 strain (**Figure 8C** and **8D**), the signal of the ACW-1-V5-eGFP fusion protein appeared cytosolic, with punctual accumulations in the distal region (**Figure 8C**), while it majorly localized around the septa (**Figure 8C**, red arrow) and at the hyphal periphery (**Figure 8C**, white arrow). ACW-1-V5-eGFP appeared to accumulate only in the outermost ring of septa, which was confirmed in a Z-stack projection (**Figure 8G** and **Video S1**). In the apical region, fluorescence appeared diffuse and cytosolic (**Figure 8D**).

**Figure 8.**
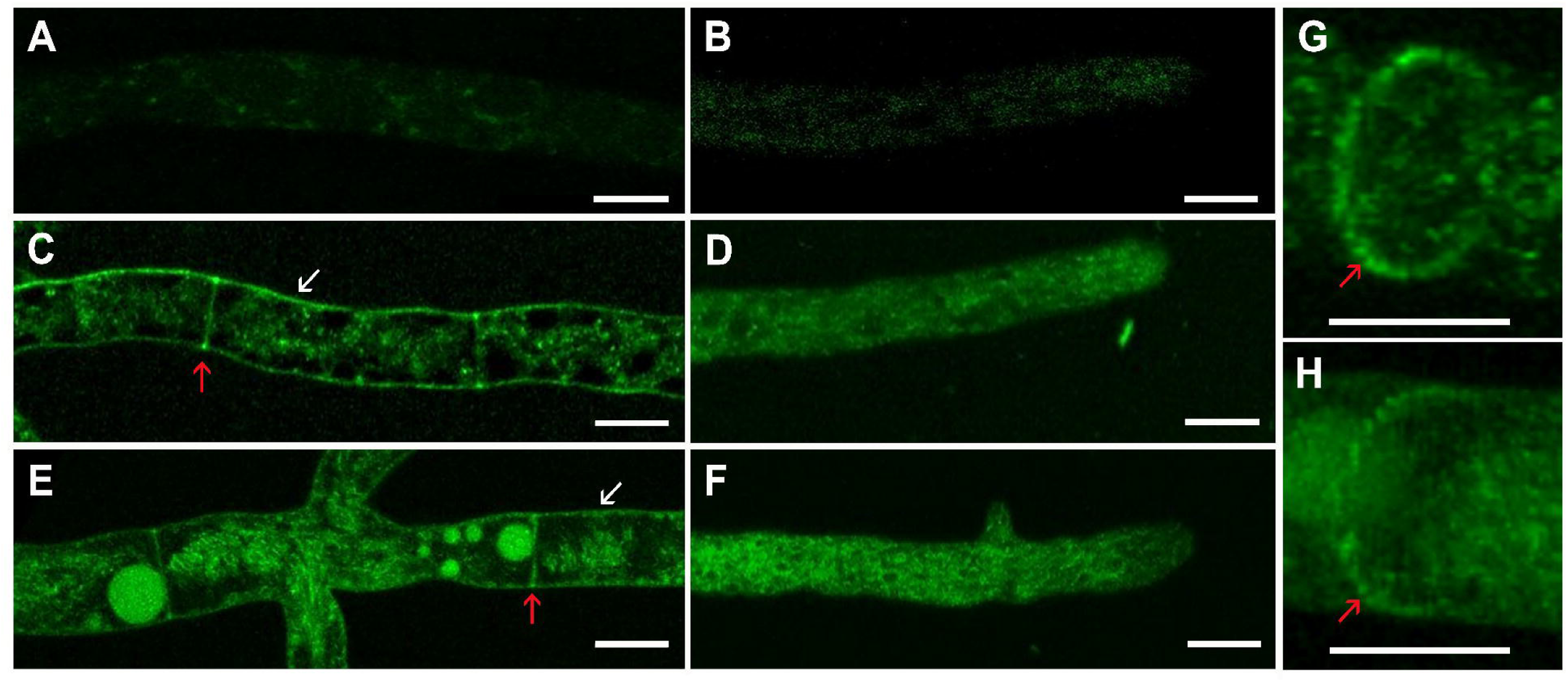
eGFP-tagged *N. crassa* ACW-1 and NCW-3 accumulate on the cell periphery and septae of mature mycelia. The parental Nc FGSC9717, Nc NCACW-1 and NCNCW-3 strains expressing eGFP-tagged ACW-1 and NCW-3 were visualized with LSCM. As expected, the parental Nc FGSC9717 strain did not generate a significant eGFP signal in the distal (A) and apical (B) regions. The Nc NCACW-1 strain produced an eGFP fluorescent signal in the periphery (C, white arrow) and septa (C, red arrow) of distal hyphae. Fluorescent signal in the apical region (D) was diffuse and appeared cytosolic. The Nc NCNCW-3 strain displayed an eGFP signal in the periphery (E, white arrow) and septa (E, red arrow) of distal regions. Apical signal (F) was diffuse and appeared cytosolic. Z-projections of Nc NCACW-1 (G) and Nc NCNCW-3 (H) strains septa showed fluorescent signal accumulation in the outermost ring (red arrow). Scale bar, 10 μm. Images were contrasted with Photoshop [131].

A similar distribution was observed in the Nc NCNCW-3 strain (**Figures 8E** and **8F**). In the distal region, the NCW-3-V5-eGFP fusion fluorescence was also localized in the outermost septum ring (**Figure 8E**, red arrow; **Video S2**), also confirmed with a Z-stack projection (**Figure H**, red arrow). Fluorescence was also detected throughout the hyphal periphery (**Figure 8E**, white arrow; **Video S2**), although a cytosolic signal was also observed. In the apical region, the signal of the NCW-3-V5-eGFP fusion seemed cytosolic and diffuse (**Figure 8F**). Unlike eGFP-V5-ACW-1, NCW-3-V5-eGFP accumulation in the cell periphery was homogeneously distributed (**Video S2**). Fluorescence signal of eGFP is known to decrease once exposed to the extracellular environment [71]. To further evaluate ACW-1 and NCW-3 surface exposure, immunofluorescence assays without permeabilization were performed with an anti-V5-Alexa 555 antibody (**Figure 9**) [23,72]. As expected, no eGFP signal was detected in the distal (**Figure 9A**) and apical regions (**Figure 9C**) of the parental Nc FGSC9717 strain. Alexa Fluor 555 did not stain the distal (**Figure 9B**) and apical (**Figure 9D**) regions, either. These results confirmed no expression of eGFP and discarded non-specific Alexa Fluor 555 binding. In contrast, the Nc NCACW-1 strain produced an eGFP signal in the septum and the periphery of distal (**Figure 9E**) regions, confirming results already shown in **Figure 8**. A cytosolic and a faint periphery signal was detected in the apical (**Figure 9H**) region. Alexa Fluor 555 signal was also detected at the periphery and septa of distal regions (**Figure 9F**), partially colocalizing with eGFP signal (**Figure 9G**). In the apical region, Alexa Fluor 555 signal was also detected along the diameter of the hyphae and periphery (stronger than eGFP signal) (**Figure 9I**), with partial co-localization with the eGFP signal (**Figure 9J** and **K**). Anti-V5-Alexa Fluor 555 immunofluorescence on Nc NCNCW-3 strain did not produce additional information besides that observed with eGFP imaging (data not shown).

**Figure 9.**
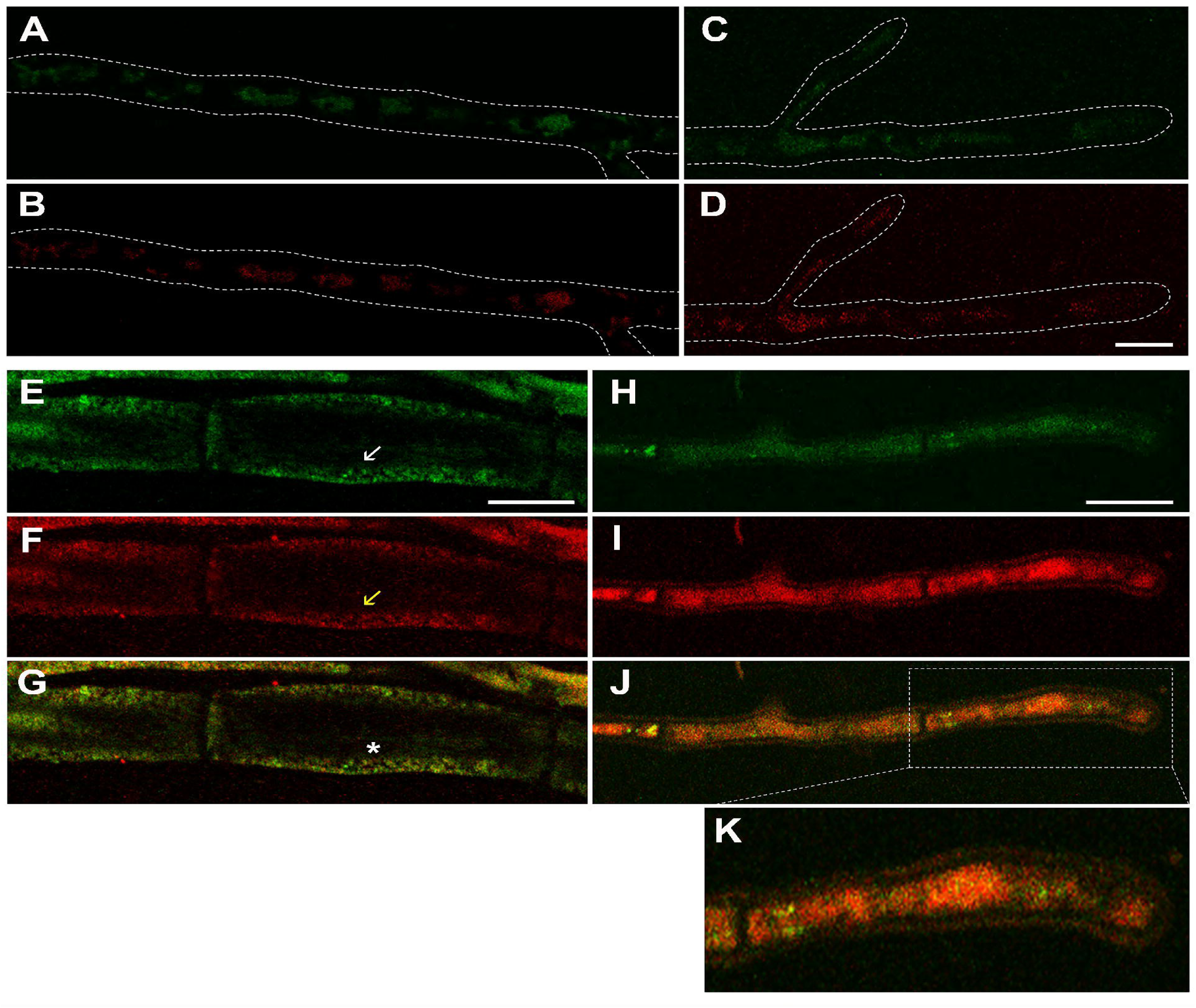
Immunofluorescence of Nc NCACW-1 strain. The parental Nc FGSC9717, and Nc NCACW-1 strains were stained with anti-V5-Alexa Fluor 555 antibodies without a permeabilizing step to further evaluate their localization. As expected, the parental Nc FGSC9717 strain did not produce any eGFP signal in the distal (A) and apical (C) regions. No Alexa Fluor 555 signal was detected in distal (B) and apical (D) regions either. The Nc NCACW-1 strain showed eGFP signal in the periphery and the septa in the distal (E) region, while near the tip (H), the signal was mostly cytosolic and faint in the periphery. Alexa Fluor 555 signal was also detected in the periphery and septa of distal regions (F), while it was peripheral in the apical region (I). eGFP and Alexa Fluor 555 signals partially colocalized in the distal (G) and faintly in the apical (J, K) regions. Scale bar, 10 um. Images were contrasted with Photoshop [131].

## DISCUSSION

Structural non-catalytic cell wall resident proteins evolve faster than cell wall and cytosolic enzymes [17,19,23,73,74]. This provides fungi a diversified cell wall proteome well suited for adaptation to changing environments. Rapid evolutionary rates have been reported for major classes of CWPs, including GPI-CWPs [17,19,74], the most abundant CWPs, which are covalently bound to cell wall β-1,3-glucans via transglycosylation of GPI-remnants, and PIR-proteins, the second most abundant CWPs, which bind the wall via transamidation of the conserved glutamine residue in PIR motifs [9–11].

ACW-1 is a classical GPI-CWP [14,28,29,34] and a major constituent of the *N. crassa* cell wall [14]. In contrast, NCW-3 lacks a GPI anchor and contains a CBM-52 domain associated with β-1,3-glucan recognition. Although both proteins are ultimately incorporated into the cell wall matrix, their modes of initial interaction differ. ACW-1 is targeted through its GPI anchor, whereas NCW-3 likely associates with β-1,3-glucan through its CBM-52.

We hypothesized that these distinct modes of interaction impose different structural and functional constraints, which in turn shape their evolutionary trajectories. To test this, we compared the diversity, structural features, evolution, cellular allocation and functional roles of ACW-1 and NCW-3.

### ACW-1 and NCW-3 share similar localization patterns and contribute to cell wall integrity

In *N. crassa*, eGFP-tagged ACW-1 and NCW-3 predominantly localized to the septa and cell surface of distal hyphal regions (**Figures 8** and **9**). In particular, both ACW-1 and NCW-3 accumulated at the outer ring of septa (**Figure 8**), where accumulation of glucan-peptide fibrils has been reported [75]. Given their association with β-1,3-glucans [31], this localization is consistent with their enrichment in cell wall regions where these polysaccharides are present [75–78]. Nevertheless, the absence of β-1,3-glucan synthetic machinery in septa [79,80] raises the question of how this polysaccharide is deposited in this subcellular localization.

The preferential accumulation of ACW-1 and NCW-3 in distal rather than apical regions (**Figures 8** and **9**) further supports the idea that both proteins may require the composition and structure of mature septa [75,76] for binding. Differences in stiffness between the cell walls of mature hyphae and growing tips have been identified in filamentous fungi, with mature hyphae being more rigid than the tips [81]. Mature hyphae and septa are characterized by more extensively cross-linked cell walls compared to the more dynamic apical regions, where active synthesis and remodelling predominate. In this context, accumulation of ACW-1 and NCW-3 at distal septa and cell walls is consistent with a role in reinforcing or maintaining the structural integrity of mature wall regions. These observations raise the idea that, in addition to β-1,3-glucan interaction, the structural and compositional characteristics of mature cell wall compartments may contribute to the evolutionary constraints shaping ACW-1 and NCW-3.

Functionally, deletion of either ACW-1 or NCW-3 encoding genes increased sensitivity to CR in *N. crassa*, indicating compromised cell wall integrity consistent with altered β-1,3-glucan organization (**Figure 7**). While the magnitude of the phenotype differed between KO mutants (see below), they support a role for both proteins in the structural organization and stabilization of the β-1,3-glucan network (**Figure 7**). All these results indicate that ACW-1 and NCW-3 localize to similar cell wall regions and contribute with the cell wall integrity, likely through interactions with β-1,3-glucans. Despite differences in these modes of interaction, both proteins appear to participate in related structural functions within the cell wall matrix.

### ACW-1 and NCW-3 share **β**-1,3-glucan association and localization but differ in functional contributions to the cell wall

Although both ACW-1 and NCW-3 associate with β-1,3-glucans, localize to similar cell wall regions and share partially overlapping functions, their functional contributions to the cell wall are not fully analogous. Deletion of ACW-1 had limited impact on the vegetative growth under standard conditions, with no major differences in hyphal morphology (**Figure 6E**), growth rate (**Figure 6G**) and branching (**Figure 6H**) compared to the Nc WT strain. In contrast, the Nc Δ*ncw-3* strain exhibited aberrant hyphal morphology, reduced growth rate and branching compared to the Nc WT strain, which indicated a more prominent role for NCW-3 in maintaining normal hyphal development. Alterations in gross colony morphology (**Figure 6B**) and reduction in conidiation (**Figure 6I**) in both mutant strains suggested defects in the cell wall beyond basic vegetative growth, with the effect on the Nc Δ*ncw-3* strain being more severe than the Nc Δ*acw-1* strain.

Variable phenotypes have been reported for ACW-1 homologues (ECM33 proteins) KO strains in other fungi. While deletion of ECM33 proteins has resulted in minimal phenotypic effects in some organisms, such as *A. fumigatus* [82–84], more severe defects in cell wall organization and development have been described for *S. cerevisiae* [32,85,86], *C. albicans* [87–89], and other filamentous species like *A. flavus* [90] and *F. oxysporum* [91]. This variability suggests that, while ACW-1/ECM33 proteins may share conserved roles in the cell wall, the extent of their functional contribution is species-dependent.

Changes in basal cell wall composition further support a structural role for ACW-1 and NCW-3. Loss of ECM33 in other fungi has been associated with changes in glucan content and protein leakage into the media [32,90,92,93]. Therefore, ECM33 may be necessary for the correct retention of other cell wall components [94]. Both the Nc Δ*acw-1* and Nc Δ*ncw-3* strains exhibited reduced β-1,3-glucan and total cell wall protein signals, while levels of chitin signal remained unchanged under standard growth conditions (**Figures 7E-G**). These decreased signals, consistent with less content of these components in the cell wall, suggested ACW-1 and NCW-3 may also be involved in the retention of other CWPs and cell wall glucans, particularly through cross-linking. Loss of ECM33 is often accompanied by increase in chitin deposition as a compensatory mechanism [32,83,86,90–92], in contrast to the Nc Δ*acw-1* and Nc Δ*ncw-3* strains where chitin signals remained unchanged. This indicates that, under normal growth conditions, the loss of ACW-1 and NCW-3 did not trigger compensatory chitin upregulation.

Consistent with a role in the cell wall integrity, both Nc Δ*acw-1* and Nc Δ*ncw-3* showed increased sensitivity to cell wall-perturbing agents, CFW (**Figure 7A**) and CR (**Figure 7B**). However, the patterns of sensitivity differed. The Nc Δ*acw-1* exhibited a stronger response to CFW despite its overall phenotype resembling that of Nc WT. On the other hand, Nc Δ*ncw-3* displayed less sensitivity to CFW and a dose-independent response to CR, in contrast with its otherwise altered phenotype relative to the Nc WT. This suggested distinct modes of interaction with the cell wall. Additional differences were observed under abiotic stress conditions. The Nc Δ*acw-1* displayed increased resistance to osmotic stress, whereas Nc Δ*ncw-3* showed greater resistance to elevated growth temperatures. The increased resistance of both KO strains to osmotic stress was unexpected given their overall sensitivity to CFW and CR. One possible explanation could be a reorganization of the polymers in the cell wall affecting its elasticity and response to osmotic shock [95]. Alternatively, the increased resistance may reflect the activation of compensatory stress response pathways. In fungi, the cell wall integrity pathway (CWI) has been reported to cross-talk with the high-osmolarity glycerol pathway (HOG) [96–98]. Disruption of cell wall components can therefore lead to activation of compensatory stress response mechanisms, enhancing tolerance to osmotic stress. These observations suggested that loss of ACW-1 or NCW-3 altered the structural organization and the composition of the cell wall in *N. crassa*. However, strain-specific and inter-strain variability in response across stressors warrants further investigation.

### ACW-1 and NCW-3 are predominantly encoded by ascomycetes but show distinct diversification patterns within Kingdom Fungi

ACW-1 and NCW-3 homologues were predominantly identified in ascomycetous fungi with limited representation in early-diverging fungal lineages such as Mucoromycota and Zoopagomycota (**Figure 1**). This distribution suggests that both proteins represent solutions that emerged primarily in fungi with cell walls enriched in glucans, including β-glucans, such as ascomycetous cell walls [99]. Nevertheless, the distribution of ACW-1 and NCW-3 differed within ascomycetous fungi (**Figure 1**). ACW-1 was broadly represented across multiple major taxonomical families (*Aspergillaceae*, *Glomerellaceae*, *Xylariaceae*, *Nectriaceae*, *Saccharomycetaceae*) (**Figure 1A**), whereas NCW-3 showed a more restricted distribution, being largely confined to Sordariales (**Figure 1B**). In addition, ACW-1 exhibited a high degree of diversification in yeasts, where multiple paralogs were identified (**Figure 3A**). This pattern suggested that ACW-1 may be subjected to lineage-specific expansion and functional diversification [100], likely driven by selective pressures associated with the wide range of ecological niches occupied by yeasts. Similar patterns of expansion and diversification have been reported for other GPI-anchored CWPs, such as adhesins [20], which are organized in different families in yeasts like *S. cerevisiae*, *C. albicans* and *C. glabrata*, consisting of highly similar sequences that confer distinct cell surface properties. In contrast, NCW-3 displayed a more homogeneous distribution within Sordariales and stronger clustering by taxonomic family (**Figures 1B** and **3B**), consistent with a more conserved and potentially lineage-specific role within this group of fungi. This restricted diversification suggested that NCW-3 may be subject to tighter functional constraints than ACW-1. However, the coexistence of ACW-1 and NCW-3 in some species, including *N. crassa* (**Figure 1C**), indicates that these proteins are not mutually exclusive and can act within a shared cellular and structural context.

### ACW-1 and NCW-3 show similar global evolutionary change, but within ACW-1 occurred heterogeneously

Evolution of both ACW-1 and NCW-3 was not parallel to that of the species they are encoded in (**Figure 2**), showing a different evolutionary pace typical of CWPs [17,19,74]. However, both ACW-1 and NCW-3 retained moderate sequence conservation, reflected by comparable Shannon entropy distributions. Furthermore, similar nonsynonymous-to-synonymous substitution rate ratios (dN/dS) (ω: ∼0.15 for both proteins) suggested purifying selection acting on both families within shared taxa in Sordariales. These results suggested that ACW-1 and NCW-3 are subjected to similar global evolutionary change despite their distinct diversification patterns and structural features.

Nevertheless, within ACW-1, evolutionary divergence was not homogeneous. ACW- 1 shows characteristics of other CWPs that facilitate rapid evolution [73] like widespread diversification in paralog families, as is the case for yeasts, and the presence of dispersed repeats, as predicted in this work in the form of LRRs (**Figure 5A**). P-distance calculation between modern LRR regions and the reconstructed ancestral LRR resulted in a p-distance gradient within ACW-1 (**Figure 5B**). Predicted LRRs close to the N- and C-termini have a higher p-distance, consistent with higher divergence, than predicted LRRs closer to the center of the protein, one of which has a high similarity to the reconstructed ancestral sequence (**Figure 5B**). This suggests that ACW-1 is composed of a primitive nucleus that may have multiplied and expanded the protein, a mechanism that has been reported for other tandem repeat-containing CWPs like adhesins and agglutinins, perhaps through DNA replication slippage, unequal cross-over, or double strand break repair [73]. Moreover, the different p-distances from the reconstructed ancestral sequence indicates LRR units have diverged at a different pace (**Figure 4** and **5**), perhaps due to different functional or structural constraints along the protein [102]. A conservation analysis with ConSurf revealed such possible constraints (**Figure 5C** and **D**) where higher conservation on LRRs repeat units close to the middle and near the C-terminus of ACW-1 was found. These repeat units interact with the *N. crassa* cell wall (besides GPI-related processing), as evidenced by the presence of asparagine residues (N155, N240) involved in the cell wall cross-linking of ACW-1 N-glycosylations to β-1,3/1,4-glucans [14,28,34].

The p-distance to the ancestral sequence and the conservation analyses captured distinct and complementary aspects of ACW-1 evolution (**Figure 5**). The p-distance analysis reflected the historical divergence of individual repeat units relative to a reconstructed ancestral sequence and is Nc ACW-1-specific, whereas the ConSurf analysis reflected sequence constraints among extant ACW-1 homologues. Therefore, similarity to the ancestral sequence may not necessarily imply strong conservation across modern proteins, as these two analyses address different evolutionary dimensions. Central repeat units (**Figure 5A** and **B**), which retained higher similarity to the ancestral sequence, may therefore reflect preservation because of lacking strong enough evolutionary pressure to change. In contrast, the higher conservation observed in C-terminal repeat units among extant homologues, despite their greater divergence from the reconstructed ancestor, is consistent with an initial stage of strong evolutionary pressure on C-terminal repeat units to adapt and integrate into the cell wall, followed by functional constraints that prevent losing that interaction in modern proteins. In NCW-3, an analogous heterogeneous accumulation of evolutionary change was not observed.

A similar global evolutionary change, but different accumulation of such change between ACW-1 and NCW-3, may be explained by local, distinct structural features. In ACW-1, evolutionary constraints may be associated with the repetitive modular architecture, GPI anchor signal and N-glycosylation sites (**Figure S1**), whereas for NCW-3 they may be associated primarily with the conservation of the CBM-52 module (**Figure S2**). Distinct functional constraints for ACW-1 and NCW-3 could also be inferred from the KO strains, with the *N. crassa* NCW-3 KO strain displaying more severe phenotypic defects, including reduced growth rate, altered hyphal branching and decreased conidiation, indicative of a stronger functional commitment compared to ACW-1 (**Figure 6**).

The divergence of ACW-1 may be further facilitated by this lower functional commitment and a resilient fold that tolerates extensive sequence variation without loss of stability, structure and function, a feature that has been described in LRR-containing proteins [48]. This scenario is further supported by the identification of remote structural homologues of Nc ACW-1 despite low sequence identity (28%) [101]. The evolutionary constraints acting on NCW-3 are less clear; however, its stronger phenotypic impact suggests a more influential role in *N. crassa* hyphae biology (**Figure 6 and 7**).

Our comparative analyses revealed that ACW-1 and NCW-3 follow different evolutionary trajectories. Both proteins were primarily identified in Ascomycota across different lineages and distinct divergence patterns. ACW-1 displayed a broader distribution, greater sequence divergence and paralogy, particularly in Saccharomycotina yeasts, consistent with a functional diversification associated with adaptation to diverse ecological niches. In contrast, NCW-3 exhibited a more restricted distribution, predominantly within Sordariales, alongside a more homogeneous divergence pattern, suggesting lineage-specific specialization.

Despite these differences, both protein families displayed similar global evolutionary change, reflected by comparable moderate sequence conservation and signatures of purifying selection within shared taxa in Sordariales that, however, accumulated differentially in ACW-1 and NCW-3. This may be explained by the distinct structural characteristics of each protein accommodating these diversification patterns, and further supported by the associated distinct functional constraints.

Although neither protein appeared essential for *N. crassa* survival, both localize to the outermost ring of the septa and lateral walls of mature hyphae, interact with β-1,3-glucans, and contribute differently to the structure and organization of the cell wall, with NCW-3 likely playing a more prominent role. These findings suggested that shared localization and substrate association do not necessarily imply full functional analogy among structural CWPs. Furthermore, beyond cellular role and lineage distribution, local structural features may also influence the evolutionary constraints of a protein and its trajectory even within the same class. This work supports a shift toward a more nuanced, CWP evolution approach. These insights may guide future studies exploring how CWP diversification contributes to cell wall dynamics, fungal adaptation and specialization, while also generating structural and functional insights with potential applications in directed evolution and protein engineering.

## MATERIALS AND METHODS

### Bioinformatics

#### In silico structural characterization of N. crassa ACW-1 and NCW-3

Protein disorder was predicted using AIUpred [45], IUPred2A (using ANCHOR2 algorithm) [46] and ESpritz [44]. Mature tertiary structures were predicted with AlphaFold2 using MMseqs2 via Colab v1.5.3 with default settings [43] and annotated with PyMOL v3 [47]. Prediction of N-glycosylation was performed with NetNGlyc v1.0 [62].

#### Search for N. crassa ACW-1 and NCW-3 fungal homologues

Amino acid sequences of *N. crassa* ACW-1 (NCU08936, UniProt Q1K6S6_NEUCR) and NCW-3 (NCU07817, UniProt Q7SBE7_NEUCR) were retrieved from UniProt [103]. The complete ACW-1 and NCW-3 sequences were queried against the non-redundant (nr) Fungi database (taxid: 4751; September 2024) using BLASTp with the BLOSUM62 matrix [104]. Homologues were defined as proteins sharing equal or greater than 30% sequence similarity over equal or greater than 50% of the length to ACW-1 and NCW-3, with an E-value threshold equal or less than 1x10^-6^. Retrieved sequences were manually curated and screened for cell wall protein-specific features. Signal peptides were predicted with SignalP 6.0 [105], and GPI addition signals were assessed by NetGPI v1.1 [106]. ACW-1 homologues were required to possess a GPI signal, whereas NCW-3 homologues were expected to lack one. Transmembrane domains were predicted with DeepTMHMM [107]. Functional domains were predicted via locally-run hmmscan from HMMER v3.4 [55] against the Pfam-A database [108]. Overall sequence conservation was estimated with Shannon entropy calculation with the Bio3D package v2.4-5 [109] in RStudio v4.4.2 [110]. Scored residue conservation and prediction of functional or structural roles were performed with ConSurf [61]. Hidden Markov Models (HMM) and consensus sequences were generated using the hmmbuild tool from HMMER v3.4 [55]. Proteins were clustered using the EFI-Enzyme Similarity Tool [42] (EFI-EST) at minimum 85 and 75% identity threshold for ACW-1 and NCW-3 homologues, respectively. Clusters were visualized in Cytoscape 3.10.3 [111].

#### Construction of phylogenetic trees

Homologous sequences were aligned using the Multiple Alignment using Fast Fourier Transform (MAFFT) algorithm [112] through the Galaxy server [113] and manually inspected in Jalview v2 [114]. Phylogenetic trees were inferred with the Randomized Accelerated Maximum Likelihood method (RAxML) under the maximum-likelihood algorithm (ML) and automatic iterations [115]. Reconstruction was performed using the PROTGAMMAIWAG model, which applied the Whelan and Goldman substitution matrix (WAG) with a gamma distribution of rate heterogeneity. In the case of the ACW-1 homologues, rate heterogeneity included a proportion of invariant sites and empirical amino acid frequencies estimated from the alignment. For the NCW-3 homologues, invariant sites were included with fixed model frequencies. In both cases, 1000 rapid bootstrap replicates were conducted followed by a thorough maximum-likelihood search, and the autoMRE criterion was applied to automatically determine bootstrap convergence. Additional trees were generated to examine the distribution of ACW-1 and NCW-3 orthologues across fungi. For this, 4171 fungal reference genomes were retrieved from RefSeq NCBI Dataset [40] (September 2024) and organized using the NCBI taxonomy common tree tool. Trees were annotated and edited in iTOL v7.2.2 [116].

#### Estimation of non-synonymous-to-synonymous rate ratios (dN/dS)

Complete protein sequences from ACW-1 and NCW-3 homologues belonging to shared taxa were retrieved from UniProt [103]. Corresponding coding DNA sequences (CDS) were obtained from the RefSeq NCBI database [40]. Protein sequences were aligned with MAFFT [112] through the Galaxy server [113], inspected in Jalview v2 [114] and used to generate codon alignments with the CDS sequences with PAL2NAL [117]. Phylogenetic trees were inferred from the codon alignments using IQ-TREE v3 [58] and ModelFinder to select the best-fit solution model [59]. Reconstruction of the ACW-3 codon alignment tree was performed using the KOSI07+F+I+R7 model, while the reconstruction of the NCW-3 codon alignment tree used the KOSI07+F+R5 model. These models incorporated empirical codon frequencies and FreeRate heterogeneity across sites, while the ACW-1 model additionally accounted for a proportion of invariant sides. Estimation of dN/dS was performed as previously described [118,119] using the M0 codon substitution model implemented in codeml from the Phylogenetic Analysis by Maximum Likelihood (PAML) package [41].

#### LRR domain identification, ancestral LRR sequence reconstruction and evolutionary distance between LRR domains

LRR domains across ACW-1 and its homologues were identified with locally-run hmmscan from HMMER v3.4 [55] against the Pfam-A database [108]. Predicted LRR hits of a length of 20-40 amino acids and a conditional E-value < 0.05 were extracted and aligned with MAFFT [112] through the Galaxy server [113]. Redundant sequences were clustered at 95% identity with locally-run CD-HIT v4.8.1 [120]. Poorly aligned positions were trimmed with locally-run trimAI v1.5.0, using a gap threshold of 0.3, removing columns with more than 70% gaps [121]. Phylogenetic trees were inferred in IQ-TREE v3 [58] using ModelFinder [59] to select the best-fit model, and support was assessed with 1000 ultrafast bootstrap replicates [60]. Reconstruction was performed with the Q.YEAST+R7 substitution model, which utilized an empirical amino acid substitution matrix optimized for yeast proteins combined with seven discrete rate categories for across-site heterogeneity. The resulting alignments and phylogenetic trees were used as input for ancestral sequence reconstruction in IQ-TREE using empirical Bayes methods. The evolutionary distances between the reconstructed ancestral sequence and modern LRR repeat units were calculated with the p-distance method, considering gaps to account for indel events, with the SeqinR package v4.2-36 [122] in RStudio v4.4.2 [110].

### Strains, media and culture conditions

Bacterial and fungal strains used in this work are listed in **Table 1**. Bacteria were grown in Luria-Bertani (LB) [123] medium supplemented with 0.1 mg/mL ampicillin and 15 g/L agar, if necessary, and grown at 37 °C. Liquid media cultures were incubated at 220 rpm.

**Table 1.**
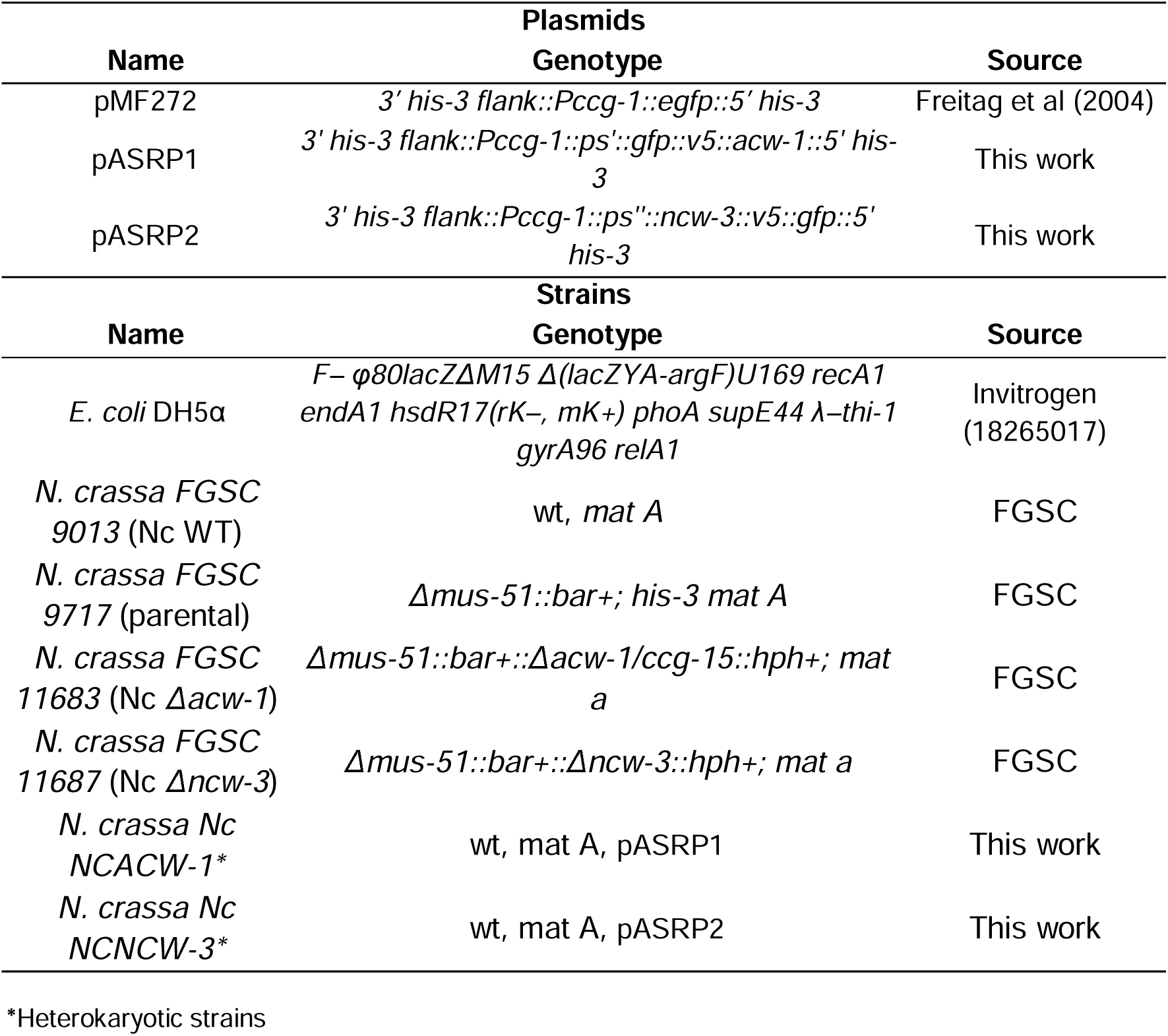
Plasmids and strains used in this work.

Fungal strains were grown and maintained on Vogel’s Minimal Medium (VMM) [124], supplemented with 15 g/L sucrose, 2 mg/mL L-histidine for auxotrophic strains, and 15 g/L agar for solid media. For the Calcofluor (CFW) and Congo Red (CR) susceptibility assays, the pH of the VMM was previously adjusted to pH 5.95 with 100 mM MES-NaOH. Transformant fungal strains were selected on FGS agar [125] without supplements. Mycelium for cell homogenates and immunofluorescence assays was cultured in complete Vogel Medium (CVM) (VMM supplemented with 0.5% w/v yeast extract and 0.5% w/v casamino acids). Fungi were cultured at 25 °C, unless otherwise specified.

### Molecular methods

#### Construction of eGFP-tagged ACW-1/NCW-3 molecular assemblies, NCACW-1 and NCNCW-3

Construction of molecular cassettes for expression of eGFP-tagged ACW-1 (NCACW-1, *Pccg-1::ps acw-1::egfp::v5::acw-1*) and NCW-3 (NCNCW-3, *Pccg-1::ncw-3::v5::egfp*) is illustrated in **Figure S5**. All DNA fragments were PCR amplified using Platinum™ Taq DNA Polymerase High Fidelity (Thermo Fisher, Waltham, MA, USA). The primers used in this work are listed in **Table S4** (synthesized by T4 DNA Oligo,Irapuato, GTO, Mexico) and were designed to include overlapping side regions to assemble the molecular cassettes. The individual *acw-1* signal peptide sequence (*acw-1 ps,* primers ACW1-2/ACW1-3), the *acw-1* gene without its native signal peptide but including its native stop codon (*acw-1*, primers ACW1-8/ACW1-9) and the *ncw-3* with its native signal peptide and without its stop codon (*ncw-3*, primers XbaI-NCW3-Fp/NCW3-4) were amplified from *N. crassa* FGSC-9013 (Nc WT) genomic DNA. The e*gfp* gene was amplified from pMF272 [126] without its starting and stop codons with primer pair ACW1-4/ACW1-5Bis and without its starting but including its stop codons with primer pair NCW3-7/BstBI-NCW3-Rp for NCACW-1 and NCNCW-3, respectively. The V5 epitope was introduced as the primer pairs ACW1-6Bis/ACW1-7Bis and NCW3-5/NCW3-6 for NCACW-1 and NCNCW-3, respectively.

All fragments were purified by 1% agarose gel electrophoresis and GenElute™Gel Extraction Kit (Sigma-Aldrich, St. Louis, MO, USA). DNA was quantified spectrophotometrically at 260 nm with a NanoDrop™ 2000/2000c (Thermo-Fisher). Fragments were then assembled together using the Gibson assembly method (Gibson et al., 2009) using the NEBuilder® HiFi DNA Assembly Master Mix (New England Biolabs, Ipswich, MA, USA). Due to their small sizes, V5 encoding sequences were integrated into the assemblies using an equimolar mixture (0.2 pmol total) of the adequate primer pairs, previously denatured at 90 °C for 1 minute and cooled down. The mixture was then added to the assembly reaction and incubated at 50 °C for 1 h.

The complete NCACW-1 assembly was amplified with primer pair FpACW-1/RpACW-1, that added side complementary regions that would be used to clone the assembly between the *ccg-1* promoter (*Pccg-1*) and the *3’ his-3* regions of the XbaI (Invitrogen, Waltham, MA, USA) and BstBI (New England Biolabs) digested pMF272 vector by Gibson assembly.

The complete NCNCW-3 cassette was amplified with the primer pair XbaI-NCW3-Fp/BstBI-NCW3-Rp, that added side XbaI and BstBI restriction sites, respectively. The assembly was ligated into pMF272 between XbaI and BstBI sites with T4 DNA ligase (Promega, Madison, WI, USA). The resulting pMF272:NCACW-1 (pASRP1) and pMF272:NCNCW-3 (pASRP2) were transformed into *E. coli* DH5α by electroporation. Transformant colonies were screened by colony PCR with primers UniNotI-pMF272-Fp/UniBstBI-pMF272-Rp, which amplified the complete constructs. Constructions were purified using the GenElute™ Plasmid Miniprep Kit (Sigma) and characterized by Sanger sequencing (LANGEBIO-CINVESTAV, Irapuato, GTO, Mexico).

#### Generation and characterization of N. crassa transformant strains

pASRP1 or pASRP2 plasmids were electroporated into the parental Nc FGSC 9717 strain conidia with an electric pulse of 7.5 kV/cm, 25 uF, and 600 ohms in a Gene Pulser XCell Total Electroporation System (Bio-Rad, Hercules, CA, USA). 1 mL of ice-cold 1 M sorbitol was added to recover the conidial suspension and kept on ice for 1 h. Transformants were plated on FGS agar without supplements and kept at 30 °C in darkness for 3-5 days until isolated white colonies appeared. Conidial sorbitol stocks were generated and kept at -16 °C for further use. Molecular characterization of the transformant strains is described in the **Supplementary Materials** and shown in **Figure S6**.

### Fungal phenotype characterization

Vegetative growth of the Nc WT (FGSC 9013) and Nc KO strains, Nc Δ*acw-1* (FGSC 11683) and Nc Δ*ncw-3* (FGSC 11687), was evaluated by inoculating 5x10^3^ conidia of each strain on VMM agar and incubated 16 h in darkness. Growth was measured for 8 h, every 2 h. Experiments were performed by quintuplicate. Gross colony morphology was visualized with a Leica EZ4 stereomicroscope (Leica Microsystems). Images were acquired with the LAS EZ software (Leica Microsystems) and edited with Fiji [127]. Hyphae diameter, number of branches and branch length were quantified across a length of 500 μm in n = 30 main hyphae with Fiji [127]. For the conidiation assay an initial inoculum of 5x10^3^ conidia was grown on VMM agar slants at 25 °C for 4 days in darkness and another 3 days at 16 h/8 h light/darkness cycles. Conidia were harvested in sterile distilled water (dH_2_O) and quantified with a hemocytometer [27]. Molecular characterization of the Nc KO strains is described in the **Supplementary Material** and shown in **Figure S4**.

#### Aniline blue adsorption assay

To examine the β-1,3-glucan present in the cell walls of the Nc WT (FGSC 9013), Nc Δ*acw-1* (FGSC 11683) and Nc Δ*ncw-3* (FGSC 11687) strains, the ability of their cell walls to bind aniline was evaluated [31]. Increasing amounts of cell wall material (0.4, 0.8, 1.2, 1.6 mg) were aliquoted, centrifuged at 3000 x *g* for 10 min and resuspended in 1 mL of 0.002% aniline blue (HYCEL). Suspensions were agitated at room temperature for 24 h, followed by centrifugation at 3000 x *g* for 10 min. The amount of unbound aniline blue was quantified with a spectrophotometer at 595 nm.

#### Coomassie brilliant blue adsorption assay

To examine the total protein content in the cell walls of the Nc WT (FGSC 9013), Nc Δ*acw-1* (FGSC 11683) and Nc Δ*ncw-3* (FGSC 11687) strains, the ability of their cell walls to bind Coomassie brilliant blue dye was evaluated [14]. Increasing amounts of cell wall material (0.1, 0.2, 0.3, 0.4, 0.5 mg) were aliquoted. dH_2_O was added to reach 0.5 mL. 0.5 mL of 1X Coomassie brilliant blue dye (Sigma) was added and mixed. The suspensions were incubated at room temperature for 30 min and gentle agitation, followed by centrifugation at 3000 x *g* for 10 min. Unbound Coomassie brilliant blue dye was quantified with a spectrophotometer at 465 nm.

#### Calcofluor white adsorption assay

To examine the chitin content in the cell walls of the Nc WT (FGSC 9013), Nc Δ*acw-1* (FGSC 11683) and Nc Δ*ncw-3* (FGSC 11687) strains, the ability of their cell walls to bind Calcofluor white (CFW) was measured. Increasing amounts of cell wall material (0.8, 1.6, 2.4, 3.2 and 4.0 mg) were aliquoted, centrifuged at 3000 x *g* for 10 min and resuspended in 1 mL of 0.15 μM CFW (Sigma-Aldrich, 910090; diluted in PBS, pH 7.4). The suspensions were incubated at room temperature and gentle agitation for 10 min and covered from light, followed by centrifugation at 3000 x *g* for 10 min. Unbound CFW was quantified in a dark 96 wells microplate (Costar, 3991) using a fluorimeter (Tecan Infinite 200Pro, 1103002481) at 370/432 nm (excitation/emission) wavelengths.

### Cell wall stress susceptibility assays

The effect of Congo Red (CR, 0-0.25 mg/mL), Calcofluor White (CFW, 0-1 mg/mL), osmotic stress (NaCl, 0-0.75 M) and temperature stress (25, 37 and 42 °C) on the Nc WT (FGSC 9013), Nc Δ*acw-1* (FGSC 11683) and Nc Δ*ncw-3* (FGSC 11687) strains was evaluated [65]. 5x10^3^ conidia of Nc WT (FGSC 9013), Nc Δ*acw-1* (FGSC 11683) and Nc Δ*ncw-3* (FGSC 11687) strains were spotted on VMM supplemented with the stressors and incubated at 25 °C for 16 h in darkness. Thermal stress treatments were incubated at 37 °C or 42 °C, while the control was kept at 25 °C. The inhibitory effect was assayed by measuring the vegetative growth rate of the strains at all treatments for 8 h, every 2 h. Experiments were performed by quintuplicate.

### Statistical analysis

Homoscedasticity was verified by the Levene test. Normal distribution was verified by Shapiro-Wilks test. Parametric analyses were performed by one-way analysis of variance (ANOVA). Non-parametric analyses were performed by Kruskal-Wallis test with *post hoc* Dunn test, and Mann-Whitney U test for pairwise comparisons, with p-value adjustment by Benjamini-Hochberg method. Statistical significance was determined at P-value < 0.05. Statistical analyses were performed in RStudio v4.4.2 [110].

### Live imaging

Strains were visualized with a LSM 710, AxioObserver inverted confocal microscope (Zeiss) at the Centro de Investigaciones en Óptica (CIO, Leon, Mexico) using the inverted agar block technique [128]. The alpha Plan-Apochromat 63x/1.46 Oil Korr M27 objective was used. eGFP was excited at 488 nm and its emission collected at 505-605 nm.

### Immunofluorescence assay

Non-permeabilizating immunofluorescence was performed as described elsewhere [23]. Briefly, glass slides were coated with 0.01% poly-L-lysine and covered with 1 mL of CVM. Media was inoculated with 1x10^5^ conidia and incubated at 30 °C for 16 h in darkness. Mycelia was washed twice with phosphate-buffered saline (PBS) buffer (NaCl, 137 mM; KCl, 2.7 mM; Na2HPO4, 10 mM; KH2PO4: 1.8 mM, pH 7.4) and then fixed with 4% (w/v) paraformaldehyde (in PBS) for 10 min. Mycelia was then washed twice with PBS. Free aldehydes were blocked with 40 mM NH_4_Cl (in PBS) for 10 min, and then washed twice with PBS. Samples were blocked with blocking solution (1% fetal bovine serum in PBS) at room temperature for 30 min and then washed twice with PBS. The procedure was carried out in darkness from hereon. Fixed hyphae were immunostained with 1:100 V5 Tag Monoclonal Antibody coupled to Alexa Fluor 555 (Invitrogen, 2F11F7) (diluted in blocking solution) for 1 h at room temperature. Stained samples were washed three times in PBS, and mounted in Fluoromount-G mounting medium (Sigma-Aldrich, F4680). Samples were analyzed with an inverted Olympus Fluoview™ FV1000 confocal microscope (Olympus, Tokyo, Japan) equipped with a 488 nm argon laser for GFP excitation (emission collected at 505–525 nm) and a 543 nm HeNe laser for Alexa Fluor 555 excitation (emission collected at 560–660 nm). A 60X Plan Apo N oil-immersion objective (NA 1.42, Olympus) was used for image acquisition. Images were acquired and processed using FV10-ASW software (version 4.0.2.9, Olympus).

## ABBREVIATIONS

CBM: carbohydrate binding module
CDS: coding DNA sequences
CFW: Calcofluor white
CR: Congo red
CVM: complete Vogel’s medium
CWI: cell wall integrity pathway
CWP: cell wall resident glycoproteins
eGFP: enhanced green fluorescent protein
FTOL: Fungal Tree of Life
GPI: glycosylphosphatidylinositol
HMM: hidden Markov models
HOG: high-osmolarity glycerol pathway
IDR: intrinsically disordered region
IQR: interquartile range
KO: knock-out
LB: Luria-Bertani
LRR: leucine rich repeats
LSCM: laser scanning confocal microscopy
ML: maximum likelihood
Nc: *Neurospora crassa*
NMR: nuclear magnetic resonance
nr: non-redundant
PAML: Phylogenetic Analysis by Maximum Likelihood
PBS: phosphate-buffered saline
PCR: polymerase chain reaction
PIR: proteins with internal repeats
RU: leucine rich repeat units
SSN: sequence similarity network
VMM: Vogel’s minimal medium
WT: wild type

## AUTHORS CONTRIBUTIONS

ASR-P performed most experimentation and wrote the manuscript; OAC-N acquired and prepared immunofluorescence confocal images. ASR-P and JV designed experiments and analyzed results; LA-D and JV acquired and administered financial support; JV edited the manuscript and conceptualized research. All authors critically reviewed and commented on the manuscript, and approved the final version.

## ACKNOWLEDGEMENTS

The authors would like to thank Mauricio Flores Moreno from CIO, León, Mexico, for kindly providing access to the LSCM. We also thank Meritxell Riquelme and Diego Delgado for kindly hosting Ana Sofía Ramírez Pelayo during her stay at the Department of Microbiology, CICESE, in Ensenada, Mexico. We appreciate the lab management work of Flor García Niño and Citlalli Rosales, Esteban Moreno for IT assistance, and Álvaro Zárate and CIATEJ staff for administrative support.

This work was supported by SENER-CONACYT, grant 245750 (LA-D and JV), and CONAHCYT-Ciencia de Frontera, grant 2019-552259 (JV). Ana Sofía Ramírez-Pelayo received MSc and PhD scholarships from SECIHTI-Mexico (CVU 004277).

## DATA AVAILABILITY STATEMENT

All data that support the findings of this study are available in the Materials and Methods, and Results sections, as well as in the Supplementary Material. Raw data is available upon request.

## Supplementary material

Materials and methods

**Table S1.** Distribution of ACW-1 orthologs in fungi.

**Table S2.** Distribution of NCW-3 orthologs in fungi.

**Table S3.** LRR domains predicted across ACW-1 and its homologues.

**Table S4.** Primers used in this work

**Figure S1. Sequence and structural conservation of ACW-1 homologues.**

**Figure S2. Sequence and structural conservation of NCW-3 homologues.**

**Figure S3. Prediction of LRR domains and reconstruction of the ancestral LRR unit sequence.**

**Figure S4. Molecular confirmation of KO *N. crassa* strains.**

**Figure S5. Molecular design and construction of eGFP::V5::ACW-1 and NCW-3::V5::eGFP expression assemblies, *pMF272::NCACW-1* and *pMF272::NCNCW-3,* respectively.**

**Figure S6. Molecular characterization of constructed heterokaryotic *N. crassa* strains.**

**Figure S7**. **Expression of eGFP-tagged ACW-1 and NCW-3 fusions.**

**Video S1. Z-stack reconstruction of *Neurospora crassa* NcNCACW-1 (*Pccg-1::ps acw-1::egfp::v5::acw-1*)**.

**Video S2. Z-stack reconstruction of *Neurospora crassa* NcNCNCW-3 (*Pccg-1::ncw-3::v5::egfp*)**.

