## Supplementary Material for "On the evolution, function and cellular fate of *Neurospora crassa* ACW-1 and NCW-3, proteins with different cell wall interaction mechanism"

**RUNNING TITLE**: *N. crassa* ACW-1 and NCW-3 cell wall proteins

Ana Sofía Ramírez-Pelayo[1], Olga A. Callejas-Negrete[2], Lorena Amaya-Delgado[1] and Jorge Verdín[1,*]

[1]Biotecnología Industrial, CIATEJ-Centro de Investigación y Asistencia en Tecnología y Diseño del Estado de Jalisco, Zapopan, JAL, Mexico

[2]Departamento de Microbiología, CICESE-Centro de Investigación Científica y de Educación Superior de Ensenada, Ensenada, BC, Mexico.

### Supplementary Material

#### Materials and methods

*Molecular characterization of N. crassa CWP KO mutants.* Nc WT (FGSC 9013, wt, mat A), Nc Δ*acw-1* (FGSC 11683, *acw-1/ccg-15*, mat a) and Nc Δ*ncw-3* (FGSC 11687, *ncw-3*, mat a) were grown for 72 hours in static liquid VMM, darkness and 25 °C. Mycelia was filtered through filter paper (Whatman, Maidstone, UK) under vacuum and rinsed twice with sterile distilled water. Harvested mycelia was frozen with liquid $N_2$, lyophilized at -80 °C and 5 mTor overnight, and then grinded to a powder with a pestle and mortar. Genomic DNA (gDNA) was extracted from freeze-dried mycelia using the GenElute Plant Genomic

DNA Miniprep Kit (Sigma, St. Louis, MO, USA) as per manufacturer's instructions, using molecular grade water as the eluting solution. The extracted gDNA was verified by agarose gel electrophoresis and quantified spectrophotometrically. Characterization of the homokaryotic KO mutants was performed by PCR with GoTaq(R) Green Master Mix (Promega, Madison, WI, USA) using 40 ng of extracted gDNA as template. Different primer pairs were used to verify the presence of the *hph* substitution cassette in the CWP KO mutants (Park et al., 2011) and the absence of their respective *acw-1* and *ncw-3* genes for further comparison with the Nc WT strain (see **Figure S4**).

*Molecular characterization of N. crassa transformant strains*. Genomic DNA (gDNA) from conidia of the heterokaryotic, transformant Nc NCACW-1 and Nc NCNCW-3 and parental Nc (*Δmus-51::bar+::his-3,* mat A) strains was obtained as described elsewhere (Rodrigues et al., 2018) with modifications. Briefly, a loopful of freshly collected conidia, or 100 $\mu$L from sorbitol (Sigma) stocks, was resuspended in 750 uL extraction buffer (200 mM Tris-HCl pH 8.5, 250 mM NaCl, 125 mM EDTA, 0.5% SDS) and 300 mg of previously sterilized 0.4-0.6 mm diameter glass beads. The suspension was vortexed for 5 minutes at maximum speed. Then, 375 uL of ice cold 3 M sodium acetate (pH 5.5) was added, and mixed by inversion. It was then placed at -20°C for 10 minutes and centrifuged at 14000 x *g* at 4°C (all centrifugations were performed at these conditions). One volume of cold isopropanol was added to the supernatant, mixed by inversion, and incubated at -20 °C for 1 h. The DNA pellet was washed with 70% and 96% ethanol, and air dried overnight. DNA was resuspended in 100 uL molecular grade water, verified by agarose gel electrophoresis and quantified spectrophotometrically. PCR was performed with

GoTaq(R) Green Master Mix (Promega, Madison, WI, USA) using 40 ng of extracted gDNA as template. Different primer pairs that bind outside and inside of the assemblies were used to verify the complete cassette insertions in the *his-3* locus (see **Figure S5**).

*Total cell homogenates and cell walls of N. crassa transformant strains*. Strains were grown in CVM for 3 days at 25 °C, 150 rpm and darkness. Mycelia was vacuum filtered, washed with sterile water and PBS (pH 7.4), and lyophilized overnight. Freeze-dried mycelia was homogenized in a pearl mill at 30 Hz and 6 min with 5 mm steel beads. Biomass was weighted and resuspended in 25 uL/mg of dry weight of 0.05 M Tris-HCl buffer (0.05 M Tris-HCl, pH 7.8, 1 mM PMSF) and considered as total homogenate. Cell wall isolation was performed using 1 g of dry mycelia for all strains. The total homogenates suspension was centrifuged at 35,000 x *g* at 4 °C for 10 min. Pellets were resuspended in 25 uL/mg of dry weight of SDS extraction buffer (50 mM Tris-HCl, pH 7.8, 2% w/v SDS, 0.1 M Na-EDTA, 0.023 mM β-mercaptoethanol, 1 mM PMSF), boiled for 5 min and centrifuged at 23,050 x *g* at 4 °C for 10 min (subsequent centrifugations were carried out in these conditions). The SDS extraction was performed three times.

*Western blot of total cell homogenates of N. crassa transformant strains*. Cell homogenates proteins were separated in a 15% SDS-PAGE (Laemmli, 1970) at 90 V for 2 h. Proteins were transferred to a nitrocellulose membrane at 10 V for 1 h. The membrane was blocked with 5% non-fat dry milk in TBST (20 mM Tris-base, 150 mM NaCl, 0.1% Tween 20, pH 7.6) at room temperature for 2 h and gentle agitation. Then, it was washed three times with TBST (pH 7.6). The membrane was blotted with 1:4000

Anti-V5-HRP monoclonal antibody (Invitrogen), in 1% non-fat dry milk in TBST (pH 7.6) at 4 °C overnight and without agitation. It was washed three times with TBST (pH 7.6). The membrane was developed using SuperSignal™ West Pico PLUS Chemiluminescent Substrate (Thermo Fisher Scientific) as per manufacturer's instructions. The membrane was visualized with a C-DiGit(R) Blot Scanner (Li-Cor).

**Table S1. Distribution of ACW-1 homologues in fungi**

| Order | Family | Homologue count |
|---|---|---|
| Eurotiales | Aspergillaceae | 106 |
| Hypocreales | Nectriaceae | 78 |
| Xylariales | Xylariaceae | 51 |
| Glomerellales | Glomerellaceae | 45 |
| Saccharomycetales | Saccharomycetaceae | 45 |
| Xylariales | Hypoxylaceae | 34 |
| Helotiales | Sclerotiniaceae | 25 |
| Chaetothyriales | Herpotrichiellaceae | 23 |
| Pleosporales | Pleosporaceae | 22 |
| Sordariales | Chaetomiaceae | 21 |
| Hypocreales | Hypocreaceae | 19 |
| Serinales | Debaryomycetaceae | 19 |
| Pichiales | Pichiaceae | 18 |
| Sordariales | Podosporaceae | 16 |
| Lipomycetales | Lipomycetaceae | 15 |
| Orbiliales | Orbiliaceae | 14 |
| Mycosphaerellales | Mycosphaerellaceae | 13 |
| Dothideales | Saccotheciaceae | 12 |
| Pleosporales | Didymellaceae | 12 |
| Thelebolales | Thelebolaceae | 12 |
| Pleosporales | Phaeosphaeriaceae | 11 |

| | | |
|---|---|---|
| Sordariales | Lasiosphaeriaceae | 11 |
| Teloschistales | Teloschistaceae | 11 |
| Onygenales | Arthrodermataceae | 10 |
| Sordariales | Sordariaceae | 10 |
| Xylariales | Sporocadaceae | 10 |
| Hypocreales | Bionectriaceae | 9 |
| Hypocreales | Cordycipitaceae | 9 |
| Mortierellales | Mortierellaceae | 9 |
| Mycosphaerellales | Teratosphaeriaceae | 9 |
| Pleosporales | Didymosphaeriaceae | 9 |
| Botryosphaeriales | Botryosphaeriaceae | 8 |
| Erysiphales | Erysiphaceae | 8 |
| Helotiales | Ploettnerulaceae | 8 |
| incertae sedis | incertae sedis | 8 |
| Onygenales | Ajellomycetaceae | 8 |
| Trapeliales | Trapeliaceae | 8 |
| Xylariales | incertae sedis | 8 |
| Diaporthales | Diaporthaceae | 7 |
| Xylariales | Apiosporaceae | 7 |
| Pezizales | Pyronemataceae | 6 |
| Botryosphaeriales | Phyllostictaceae | 5 |
| Glomerellales | Plectosphaerellaceae | 5 |
| Helotiales | Lachnaceae | 5 |
| Myriangiales | Elsinoaceae | 5 |
| Pezizales | Pezizaceae | 5 |

| | | |
|---|---|---|
| Pezizales | Tuberaceae | 5 |
| Saccharomycodales | Saccharomycodaceae | 5 |
| Coniochaetales | Coniochaetaceae | 4 |
| Diaporthales | Valsaceae | 4 |
| Dipodascales | incertae sedis | 4 |
| Helotiales | Helotiaceae | 4 |
| Lecanorales | Parmeliaceae | 4 |
| Magnaporthales | Pyriculariaceae | 4 |
| Microascales | Microascaceae | 4 |
| Onygenales | Onygenaceae | 4 |
| Pezizales | Morchellaceae | 4 |
| Pleosporales | Cucurbitariaceae | 4 |
| Pleosporales | Leptosphaeriaceae | 4 |
| Acarosporales | Acarosporaceae | 3 |
| Candelariales | Candelariaceae | 3 |
| Chaetothyriales | Trichomeriaceae | 3 |
| Cladosporiales | Cladosporiaceae | 3 |
| Helotiales | Drepanopezizaceae | 3 |
| Helotiales | Hyaloscyphaceae | 3 |
| Helotiales | Mollisiaceae | 3 |
| Hypocreales | Stachybotryaceae | 3 |
| Lichinales | Lichinaceae | 3 |
| Mycosphaerellales | Extremaceae | 3 |
| Onygenales | incertae sedis | 3 |
| Ophiostomatales | Ophiostomataceae | 3 |

| | | |
|---|---|---|
| Peltigerales | Lobariaceae | 3 |
| Phaffomycetales | Wickerhamomycetaceae | 3 |
| Serinales | Metschnikowiaceae | 3 |
| Sordariales | Schizotheciaceae | 3 |
| Venturiales | Venturiaceae | 3 |
| Xylariales | Microdochiaceae | 3 |
| Xylariales | unclassified | 3 |
| Caliciales | Physciaceae | 2 |
| Chaetothyriales | Cyphellophoraceae | 2 |
| Coniocybales | Coniocybaceae | 2 |
| Diaporthales | Gnomoniaceae | 2 |
| Dipodascales | Dipodascaceae | 2 |
| Dipodascales | Trichomonascaceae | 2 |
| Dothideales | Dothioraceae | 2 |
| Eurotiales | Trichocomaceae | 2 |
| Geoglossales | Geoglossaceae | 2 |
| Helotiales | Dermateaceae | 2 |
| Helotiales | Leptodontidiaceae | 2 |
| Helotiales | Rutstroemiaceae | 2 |
| Helotiales | Tricladiaceae | 2 |
| Helotiales | incertae sedis | 2 |
| Hypocreales | Sarocladiaceae | 2 |
| incertae sedis | Gloniaceae | 2 |
| Lecanorales | Ramalinaceae | 2 |
| Magnaporthales | Magnaporthaceae | 2 |

| | | |
|---|---|---|
| Microascales | Ceratocystidaceae | 2 |
| Mytilinidiales | Mytilinidiaceae | 2 |
| Pertusariales | Pertusariaceae | 2 |
| Phaffomycetales | Phaffomycetaceae | 2 |
| Pleosporales | Lindgomycetaceae | 2 |
| Pleosporales | Sporormiaceae | 2 |
| Pleosporales | incertae sedis | 2 |
| Teloschistales | Caloplacoideae | 2 |
| Teloschistales | Letrouitiaceae | 2 |
| Thelocarpales | Thelocarpaceae | 2 |
| Trypetheliales | Trypetheliaceae | 2 |
| Vezdaeales | Vezdaeaceae | 2 |
| Acrospermales | Acrospermaceae | 1 |
| Alaninales | Pachysolenaceae | 1 |
| Arthoniales | Chrysotrichaceae | 1 |
| Ascoideales | Saccharomycopsidaceae | 1 |
| Aulographales | Aulographaceae | 1 |
| Aulographales | Rhizodiscinaceae | 1 |
| Botryosphaeriales | Aplosporellaceae | 1 |
| Botryosphaeriales | Saccharataceae | 1 |
| Calosphaeriales | Pleurostomataceae | 1 |
| Candelariales | Pycnoraceae | 1 |
| Capnodiales | Piedraiaceae | 1 |
| Cephalothecales | Cephalothecaceae | 1 |
| Diaporthales | Cryphonectriaceae | 1 |

| | | |
|---|---|---|
| Diaporthales | Schizoparmaceae | 1 |
| Dimargaritales | Dimargaritaceae | 1 |
| Eremomycetales | Eremomycetaceae | 1 |
| Eurotiales | Elaphomycetaceae | 1 |
| Eurotiales | Thermoascaceae | 1 |
| Harpellales | Legeriomycetaceae | 1 |
| Helotiales | Hamatocanthoscyphaceae | 1 |
| Helotiales | Hyphodiscaceae | 1 |
| Helotiales | Pezizellaceae | 1 |
| Helotiales | Pleuroascaceae | 1 |
| Hypocreales | Clavicipitaceae | 1 |
| Hypocreales | Ophiocordycipitaceae | 1 |
| Hypocreales | incertae sedis | 1 |
| incertae sedis | Myxotrichaceae | 1 |
| incertae sedis | Pseudoperisporiaceae | 1 |
| incertae sedis | Thyridiaceae | 1 |
| incertae sedis | Zopfiaceae | 1 |
| Lecanorales | Lecanorineae | 1 |
| Lecanorales | incertae sedis | 1 |
| Leotiales | Tympanidaceae | 1 |
| Lichinales | Peltulaceae | 1 |
| Lineolatales | Lineolataceae | 1 |
| Lulworthiales | Lulworthiaceae | 1 |
| Microthyriales | Microthyriaceae | 1 |
| Mycosphaerellales | Dissoconiaceae | 1 |

| | | |
|---|---|---|
| Myriangiales | Myriangiaceae | 1 |
| Mytilinidiales | Argynnaceae | 1 |
| Onygenales | Arachnomycetaceae | 1 |
| Onygenales | Ascosphaeraceae | 1 |
| Onygenales | unclassified | 1 |
| Ophiostomatales | Ophiostomataceae | 1 |
| Ostropales | Stictidaceae | 1 |
| Patellariales | Patellariaceae | 1 |
| Peltigerales | Peltigeraceae | 1 |
| Pertusariales | Icmadophilaceae | 1 |
| Pezizales | Ascobolaceae | 1 |
| Pezizales | Ascodesmidaceae | 1 |
| Pezizales | Discinaceae | 1 |
| Phaeomoniellales | Phaeomoniellaceae | 1 |
| Phaeotrichales | Phaeotrichaceae | 1 |
| Pleosporales | Amniculicolaceae | 1 |
| Pleosporales | Corynesporascaceae | 1 |
| Pleosporales | Delitschiaceae | 1 |
| Pleosporales | Diademaceae | 1 |
| Pleosporales | Dothidotthiaceae | 1 |
| Pleosporales | Lentitheciaceae | 1 |
| Pleosporales | Lophiostomataceae | 1 |
| Pleosporales | Lophiotremataceae | 1 |
| Pleosporales | Melanommataceae | 1 |
| Pleosporales | Pleomassariaceae | 1 |

| | | |
|---|---|---|
| Pleosporales | Tetraplosphaeriaceae | 1 |
| Pleosporales | Torulaceae | 1 |
| Pleosporales | Trematosphaeriaceae | 1 |
| Sareales | Zythiaceae | 1 |
| Sarrameanales | incertae sedis | 1 |
| Schizosaccharomycetales | Schizosaccharomycetaceae | 1 |
| Serinales | incertae sedis | 1 |
| Sordariales | Diplogelasinosporaceae | 1 |
| Sordariales | Naviculisporaceae | 1 |
| Sordariales | incertae sedis | 1 |
| Togniniales | Togniniaceae | 1 |
| Umbilicariales | Umbilicariaceae | 1 |
| Venturiales | Cylindrosympodiaceae | 1 |
| Venturiales | Sympoventuriaceae | 1 |
| Verrucariales | Verrucariaceae | 1 |
| Xylariales | Diatrypaceae | 1 |
| Xylariales | Pseudomassariaceae | 1 |
| Xylobotriales | Cirrosporiaceae | 1 |
| Xylonales | Xylonaceae | 1 |

**Table S2. Distribution of NCW-3 homologues in fungi**

| Order | Family | Orthologs count |
|---|---|---|
| Sordariales | Chaetomiaceae | 17 |
| Sordariales | Podosporaceae | 16 |
| Sordariales | Lasiosphaeriaceae | 11 |
| Dothideales | Saccotheciaceae | 10 |
| Sordariales | Sordariaceae | 8 |
| Hypocreales | Cordycipitaceae | 6 |
| Botryosphaeriales | Phyllostictaceae | 5 |
| Botryosphaeriales | Botryosphaeriaceae | 4 |
| Magnaporthales | Pyriculariaceae | 4 |
| Microascales | Microascaceae | 4 |
| Agaricales | Mycenaceae | 3 |
| Coniochaetales | Coniochaetaceae | 3 |
| Ophiostomatales | Ophiostomataceae | 3 |
| Sordariales | Schizotheciaceae | 3 |
| Dothideales | Dothioraceae | 2 |
| Mycosphaerellales | Mycosphaerellaceae | 2 |
| Sordariales | incertae sedis | 2 |
| Trypetheliales | Trypetheliaceae | 2 |
| Venturiales | Venturiaceae | 2 |
| Agaricales | Omphalotaceae | 1 |
| Arthoniales | Chrysotrichaceae | 1 |
| Aulographales | Aulographaceae | 1 |
| Capnodiales | Capnodiaceae | 1 |
| Capnodiales | Piedraiaceae | 1 |
| Cephalothecales | Cephalothecaceae | 1 |
| Chaetosphaeriales | Chaetosphaeriaceae | 1 |
| Coniosporales | incertae sedis | 1 |
| Dothideales | unclassified | 1 |
| Magnaporthales | Magnaporthaceae | 1 |
| Mycosphaerellales | Teratosphaeriaceae | 1 |
| Mytilinidiales | Mytilinidiaceae | 1 |
| Pertusariales | Pertusariaceae | 1 |
| Pleosporales | Lophiotremataceae | 1 |
| Pleosporales | Melanommataceae | 1 |
| Pleosporales | Pleomassariaceae | 1 |
| Sordariales | Diplogelasinosporaceae | 1 |
| Sordariales | Naviculisporaceae | 1 |
| Venturiales | Cylindrosympodiaceae | 1 |

| | | |
|---|---|---|
| incertae sedis | Gloniaceae | 1 |

**Table S3. LRR domains predicted across ACW-1 and its homologues**

| *LRR domain* | *Accession* | *Description* |
|---|---|---|
| ARM_LRRK2 | PF23744.1 | Armadillo repeat (ARM) found at the N-terminus of leucine-rich repeat Ser/Thr-protein kinase 2 in eukaryotes |
| LRR_1 | PF00560.39 | Leucine-rich repeat |
| LRR_11 | PF18831.7 | Leucine-rich repeat (one repeat unit) |
| LRR_13 | PF23286.1 | Region of leucine-rich repeat in disease resistance protein RPS4B (*A. thaliana*) |
| LRR_14 | PF23598.1 | Leucine-rich repeat region found in plant resistance proteins |
| LRR_15 | PF24969.1 | Leucine-rich repeat domain found in uncharacterized fungal proteins |
| LRR_2 | PF07723.19 | Leucine-rich repeat |
| LRR_4 | PF12799.13 | Leucine-rich repeat (two copies) |
| LRR_5 | PF13306.12 | BspA leucine-rich repeat region (six copies); BspA-like surface antigens from *Trichomonas vaginalis* |
| LRR_6 | PF13516.12 | Leucine-rich repeat |
| LRR_8 | PF13855.12 | Leucine-rich repeat |
| LRR_9 | PF14580.12 | Leucine-rich repeat |
| LRR_At1g61320_AtMIF1 | PF23622.1 | Leucine-rich repeats found in F-box/LRR-repeat containing plant proteins At1g61320 and At1g61330 (*A. thaliana*) involved in the ubiquitin-proteasome pathway |
| LRR_At5g56370 | PF24758.1 | Leucine-rich repeats found in FBD-associated F-box protein At5g56370 (*A. thaliana*) and other similar plant proteins |
| LRR_ComC | PF24141.1 | Leucine-rich repeats found at the N-terminus of EGF-like domain containing protein ComC from amoebas |
| LRR_EndoS | PF23952.1 | Leucine-rich repeat domain found in Endo-beta-N-acetylglucosaminidase EndoS (*Streptococcus spp.*) |
| LRR_FBXL18 | PF19729.6 | Leucine-rich repeats from F-box/LRR repeat protein 18, FBXL18 |
| LRR_LRWD1 | PF23211.1 | Leucine-rich repeat domain found at the N-terminus of Leucine-rich repeat and WD repeat-containing protein 1 (LRWD1) from animals and related sequences |
| LRR_NXF1-5 | PF24048.1 | Leucine-rich repeat domain found in nuclear RNA export factor 1, 2, 3 and 5, NXF1, NXF2, NXF3 and NXF5, from animals and their yeast orthologues mRNA export factor 67, MEX67 |
| LRR_R13L1-DRL21 | PF25019.1 | Leucine-rich repeats found in R13L1, DRL21 (*A. thaliana*) and related plant proteins |

| | | |
|---|---|---|
| LRR_RPS2 | PF23247.1 | Leucine-rich repeats found in disease resistance protein RPS2 (*A. thaliana*) and similar plant proteins |
| LRR_Zer-1 | PF25013.1 | Leucine-rich repeats found in Zyg eleven-related protein 1 (*C. elegans*) and similar eukaryotic proteins |
| LRR19-TM | PF15176.11 | Single-span transmembrane region of LRRC19, a leucine-rich repeat protein family that functions as a transmembrane receptor inducing pro-inflammatory cytokines |

### Table S4. Primers used in this work

| Name | Sequence (5' - 3') | General Tm (°C) |
|---|---|---|
| NCW3-3 | CATCAACCAAAATGGTGGTAGAACCGAAATC | 59 |
| NCW3-4 | GTTTGCCGGTATACTGCCAAGCACC | 59 |
| NCW3-5 | GCAGTATACCGGCAAACCGATTCCGAACCCGCTGCTG GGCCTGGATAGCACCGTGAGCAAG | 59 |
| NCW3-6 | CTTGCTCACGGTGCTATCCAGGCCCAGCAGCGGGTTC GGAATCGGTTTGCCGGTATACTGC | 59 |
| NCW3-7 | ATAGCACCGTGAGCAAGGGCGAG | 59 |
| ACW1-2 | ATCAACCAAAATGCTTGTCAAGTACCTTGC | 59 |
| ACW1-3 | TTGCTCACCGACTGGGCTATAAGAGG | 59 |
| ACW1-4 | CCAGTCGGTGAGCAAGGGCGAG | 59 |
| ACW1-5BIS | GGTTTGCCCTTGTACAGCTCGTCCATG | 59 |
| ACW1-6BIS | CTGTACAAGGGCAAACCGATTCCGAACCCGCTGCTGG GCCTGGATAGCACCGCCTCTAC | 59 |
| ACW1-7BIS | GTAGAGGCGGTGCTATCCAGGCCCAGCAGCGGGTTC GGAATCGGTTTGCCCTTGTACAG | 59 |
| ACW1-8 | ATAGCACCGCCTCTACCTGCACTG | 59 |
| ACW1-9 | CGCCCTAATTACAGAAGGAGCTGGGC | 59 |
| Fhph | CACTGACGGTGTCGTCCATC | 59 |
| hphACW1 | GGATGTGATGCCCAAGACGG | 59 |
| hphNCW3 | TGATGAACATGCTCGAGATGTCTAGG | 59 |
| HIS3-3 | ATTTTCATTTAGAAGGAGCAGTCCATCTGCG | 59 |
| HIS3-4 | GGTTTGCCCATTTTGGTTGATGTGAGGG | 59 |
| FpACW-1 | CCCTCACATCAACCAAATATGCTTGTCAAGTACCTTG C | 59 |

| | | |
|---|---|---|
| RpACW-1 | GCAACTAAACGGAAATTCTTTTTTACAGAAGGAGCTG GG | 59 |
| XbaI-NCW3-Fp | AATCTAGAATGGTGGTAGAACCGAAATCATC | 59 |
| BstBI-NCW3-Rp | TTTCGAATTACTTGTACAGCTCGTCCATG | 59 |
| UniNotI-pMF272-Fp | GCGAACGAAACCCCTGAAAC | 56 |
| UniBstBI-pMF272-Rp | TCAGCATCCGTCTTGAGC | 56 |
| hisflankF | TTGATTGACAGCGAACGAAACC | 63 |
| pMF272f | AATCAACACAACACTCAAACCAC | 63 |

**FIGURES**

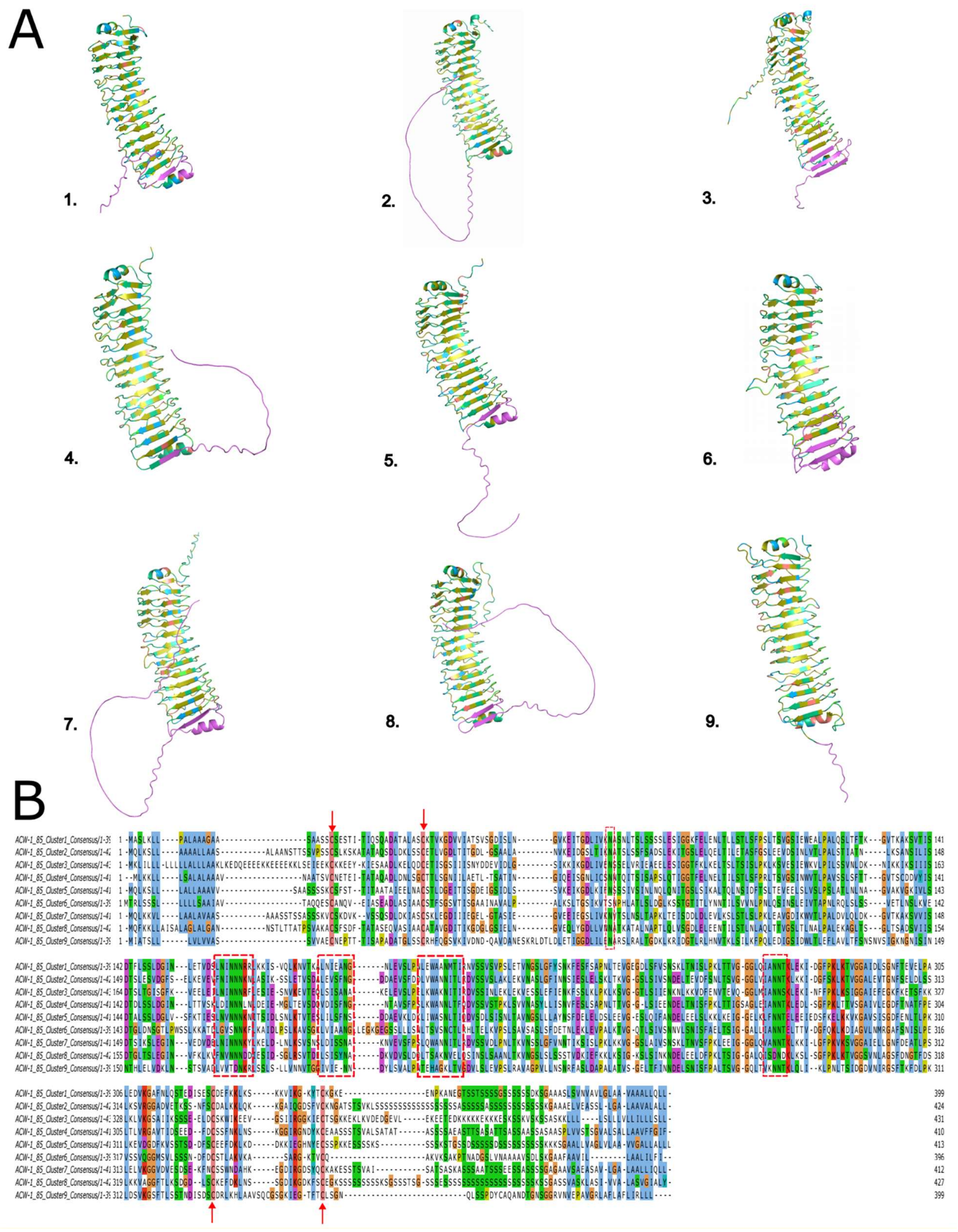

**Figure S1. Sequence and structural conservation of ACW-1 homologues.** A. Tertiary structure prediction of representative ACW-1 homologues from different clusters: 1. cluster 1: GPI-anchored cell wall organization protein Ecm33 from *Aspergillus fumigatus* Af293; 2. cluster 2: Ecm33p (YBR078W) from *Saccharomyces cerevisiae*; 3. cluster 3: Sps2p (YDR522C) from *S. cerevisiae*; 4. cluster 4: hypothetical protein V1525DRAFT_406826 from *Lipomyces kononenkoae*; 5. cluster 5: ECM33 (CAALFM_C502460CA) from *Candida albicans*; 6. cluster 6: hypothetical protein BGZ76_00099 from *Entomortierella beljakovae*; 7. cluster 7: ECM33 (YALI1_D09566g) from *Yarrowia lipolytica*; 8. cluster 8: ECM33 (Cantr_05313) from *Candida viswanathii*, and 9. cluster 9: hypothetical protein F4811DRAFT_566839) from *Daldinia bambusicola*. Retrieved ACW-1 homologues conserved the same fold formed by a mostly hydrophobic core (yellow), surrounded by polar (green) amino acids and arranged in a highly-structured β-helix. They also presented an N-terminal α-helix and a C-terminal intrinsically disordered region (IDR). Predicted structures were generated with AlphaFold2 (Jumper et al., 2021) using MMseqs2 (ColabFold v1.5.5) and visualized in PyMOL v3 (Schrödinger, LLC, 2015). B. ACW-1 homologues were clustered at a minimum 85% identity threshold. N-terminal and C-terminal cysteine residues were conserved in all clusters (pink, arrow), as well as a pair of asparagine residues near the middle of the protein (green, dotted red box) and two glycine residues on the C-terminus (orange, arrows). Hydrophobic residues were highly conserved across all proteins in different clusters (blue, arrows). The multiple sequence alignment of ACW-1 homologues clusters consensus sequences revealed highly conserved amino acid residues like N- and C-terminal cysteine residues (pink, indicated by arrows), a pair of asparagine residues in the middle of the protein (green, red dotted box) and near the C-terminus. Two glycine residues were highly conserved near the C-terminus (orange, arrows), as well as a xG motif, where x is a hydrophobic amino acid. Conserved hydrophobic residues were found spanning the alignment (blue, arrows). The sequences within each cluster were aligned with MAFFT (Katoh & Standley, 2013) and the consensus sequence was generated using hh-make from the hh-suite3 (Steinegger et al., 2019). Cluster consensus sequences were aligned with MAFFT (Katoh & Standley, 2013) and visualized with Jalview (Waterhouse et al., 2009).

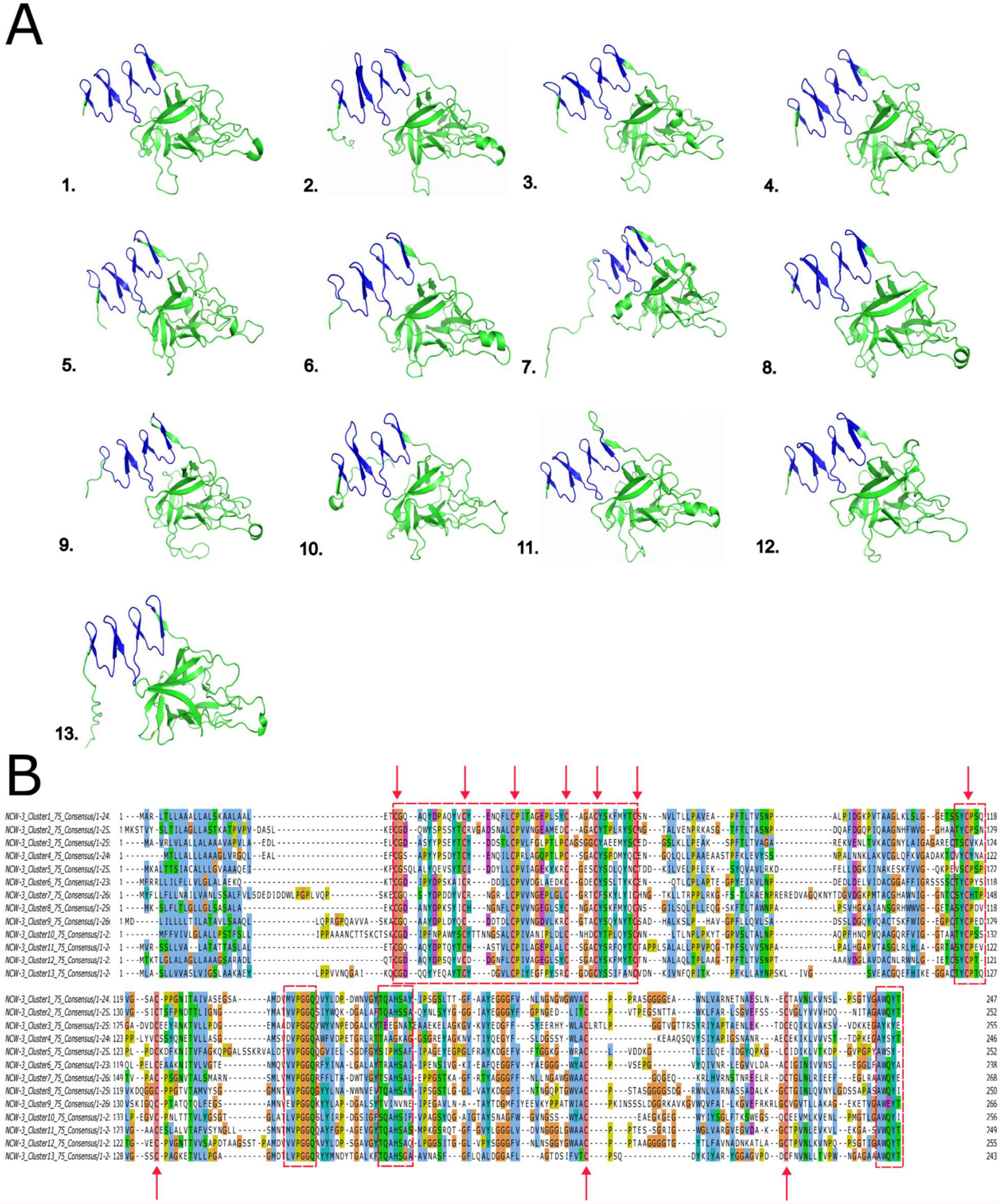


**Figure S2. Sequence and structural conservation of NCW-3 homologues.** A. Tertiary structure prediction by AlphaFold of the representative NCW-3 homologues from each cluster: 1. cluster 1: carbohydrate binding-domain-containing protein (B0T23DRAFT_313059) from *Neurospora hispaniola*; 2. cluster 2: endo-1,3(4)-beta-glucanase 1 carbohydrate binding domain-containing protein from *Aureobasidium namibiae*; 3. cluster 3: endo-1,3(4)-beta-glucanase 1 carbohydrate binding domain-containing protein (SAPIO_CDS6453) from *Pseudallescheria apiosperma*; 4. cluster 4: endo-1,3(4)-beta-glucanase 1 carbohydrate binding domain-containing protein (QBC38DRAFT_376759) from *Podospora fimiseda*; 5. cluster 5: hypothetical protein IWX90DRAFT_489274 from *Phyllosticta citrichinensis*; 6. cluster 6: secreted protein DIS24_g6347 from *Lasiodiplodia hormozganensis*; 7. cluster 7: carbohydrate-binding module family 52 protein (BBO_05441) from *Beauveria brongniartii*; 8. cluster 8: endo-1,3_4_-beta-glucanase 1 carbohydrate binding domain-containing protein (PgNI_09513) from *Pyricularia grisea;* 9. cluster 9: endo-1,3(4)-beta-glucanase 1 carbohydrate binding domain-containing protein (EV356DRAFT_500106) from *Viridothelium virens*; 10. cluster 10: carbohydrate-binding module family 52 protein (TI39_contig53g00004) from *Zymoseptoria brevis*; 11. cluster 11: carbohydrate-binding-domain-containing protein from *Schizothecium conicum*, 12. cluster 12: endo-1,3_4_-beta-glucanase 1 carbohydrate binding domain-containing protein (SPSK_06572) from *Sporothrix schenckii,* and 13. cluster 13: carbohydrate-binding module family 52 protein from *Venturia nashicola*. Most NCW-3 representatives of each cluster conserved the bilobal structure, composed of a N-terminal β-hairpin group and a C-terminal β-barrels. Predicted structures were generated with AlphaFold2 (Jumper et al., 2021) using MMseqs2 (ColabFold v1.5.5) and visualized in PyMOL v3 (Schrödinger, LLC, 2015). B. The multiple sequence alignment of NCW-3 homologues clusters consensus sequences revealed a highly conserved N-terminal amino acid region, covering the predicted β-hairpin group, by the presence of motifs such as CG, LCPx, ACYS and YxC, previously described as part of CBM-52. Four cysteine residues near the middle and C-terminus of the sequences were also conserved (pink, arrows) as well as a VPGGQ motif (red dotted box). The sequences within each cluster were aligned with MAFFT (Katoh & Standley, 2013) and the consensus sequence was generated using hh-make from the hh-suite3 (Steinegger et al., 2019). Cluster consensus sequences were aligned with MAFFT (Katoh & Standley, 2013) and visualized with Jalview (Waterhouse et al., 2009).

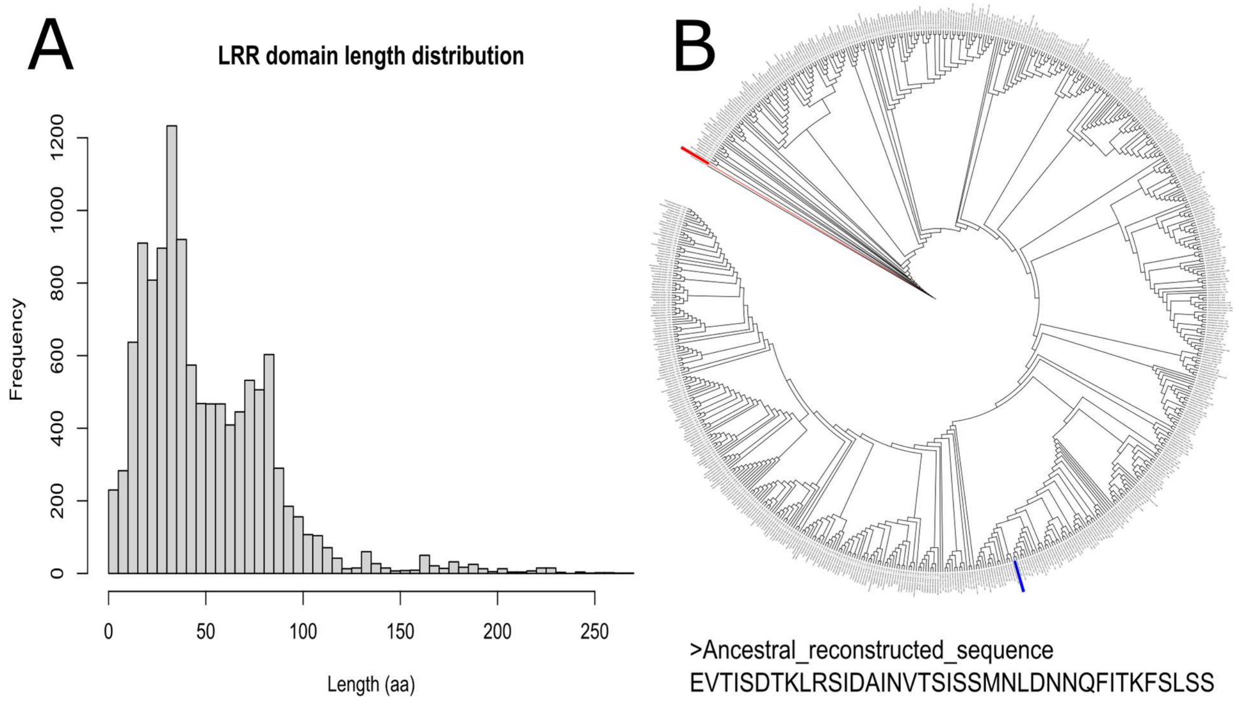


**Figure S3. Prediction of LRR domains and reconstruction of the ancestral LRR unit sequence.** A) Predicted LRR domains varied in length, while most spanned 35 residues. B) Phylogenetic tree of the ancestral reconstructed LRR sequence (bottom). Nc LRR1 (red) showed the greatest similarity to the ancestral sequence, while Nc LRR3 (blue) was more divergent and located farther in the tree. LRR domains were predicted with hmmscan in HMMER v3.4 (Potter et al., 2018) against the Pfam-A database (Mistry et al., 2021). The frequency histogram was generaled with the ggplot2 package in RStudio (Wickham, 2016). The ancestral reconstructed tree was generated in IQTREE (Hoang et al., 2018; Kalyaanamoorthy et al., 2017; Minh et al., 2020).

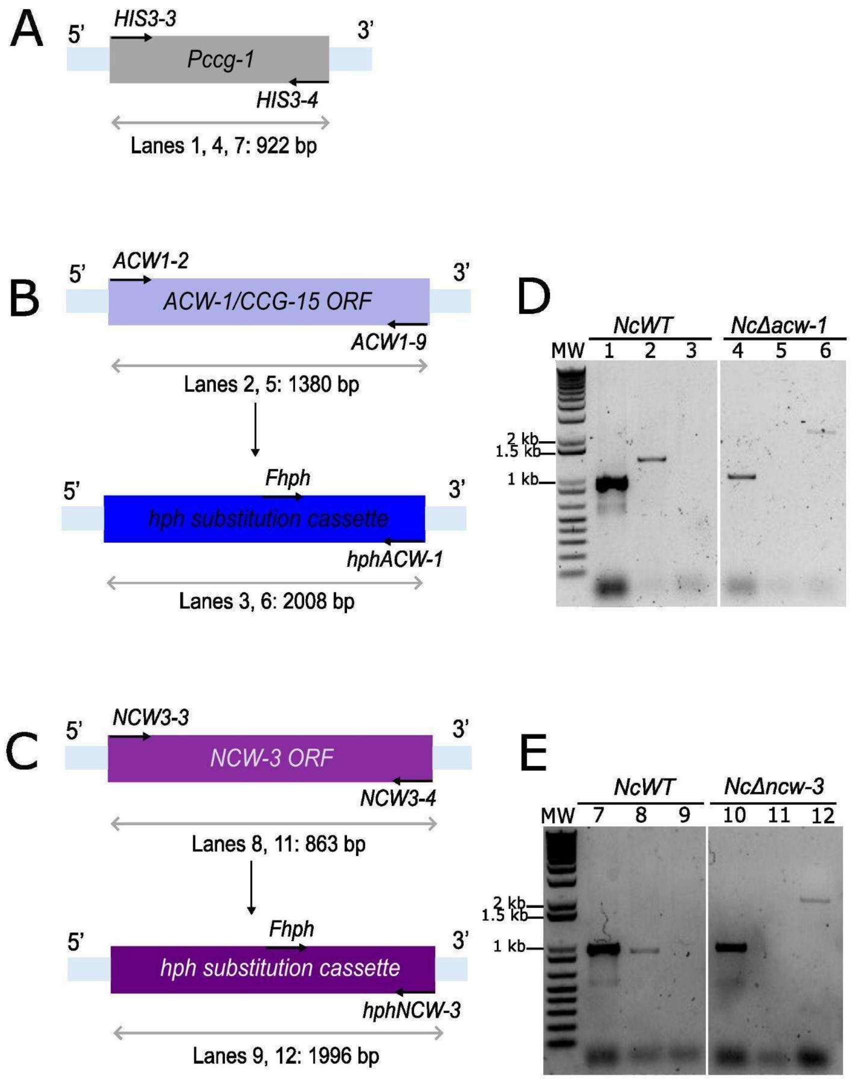


**Figure S4. Molecular confirmation of KO *N. crassa* strains.** A, D and E) PCR amplification of the *ccg-1* promoter (*Pccg-1*) with primers HIS3-3 and HIS3-4 was used as a loading control of all characterized strains: Nc WT (FGSC 9013, wt, mat A; D, lane 1; E, lane 7), Nc Δ*acw-1* (FGSC 11683, *acw-1/ccg-15*, mat a; D, lane 4) and Nc Δ*ncw-3* (FGSC 11687, *ncw-3*, mat a; E, lane 10). A band of expected size (922 bp) was obtained for all cases as the promoter was present in the three strains. B and D) The presence of the *acw-1* gene in Nc WT and its *hph* substitution in Nc Δ*acw-1* was assessed by PCR amplification of the *acw-1* gene with primers ACW1-2 and ACW1-9 from gDNA from both strains. As anticipated, a band of expected size (1380 bp) appeared in the Nc WT strain (D, lane 2) but did not amplify from the Nc Δ*acw-1* strain (D, lane 5). On the other hand, to assess the substitution of *acw-1* by the *hph* cassette, *hph* was PCR amplified with primers Fhph and hphACW-1 using gDNA from Nc WT and Nc Δ*acw-1* as templates. As anticipated, a band of the expected size (2008 bp) was obtained for the Nc Δ*acw-1* knock-out strain (D, lane 6), while it was absent from the Nc WT control strain (D, lane 3). C and E) The presence of *ncw-3* and its *hph* substitution in Nc WT and Nc Δ*ncw-3* strains was assessed by PCR amplification of *ncw-3* with primers NCW3-3 and NCW3-4 using gDNA of Nc WT and Nc Δ*ncw-3* as templates. As anticipated, a band of the expected size (863 bp) was obtained for the Nc WT control strain (E, lane 8) and was absent from the Nc Δ*ncw-3* knock-out strain (E, lane 11). In parallel, the presence of the *hph* substitution cassette used to knock out the *ncw-3* gene was evaluated by PCR amplification with primers Fhph and hphNCW3 using gDNA from Nc WT and Nc Δ*ncw-3* strains. As anticipated, a band of expected size (1996 bp) was obtained for the Nc Δ*ncw-3* knock-out strain (E, lane 12), but was absent from the Nc WT control strain (E, lane 9).

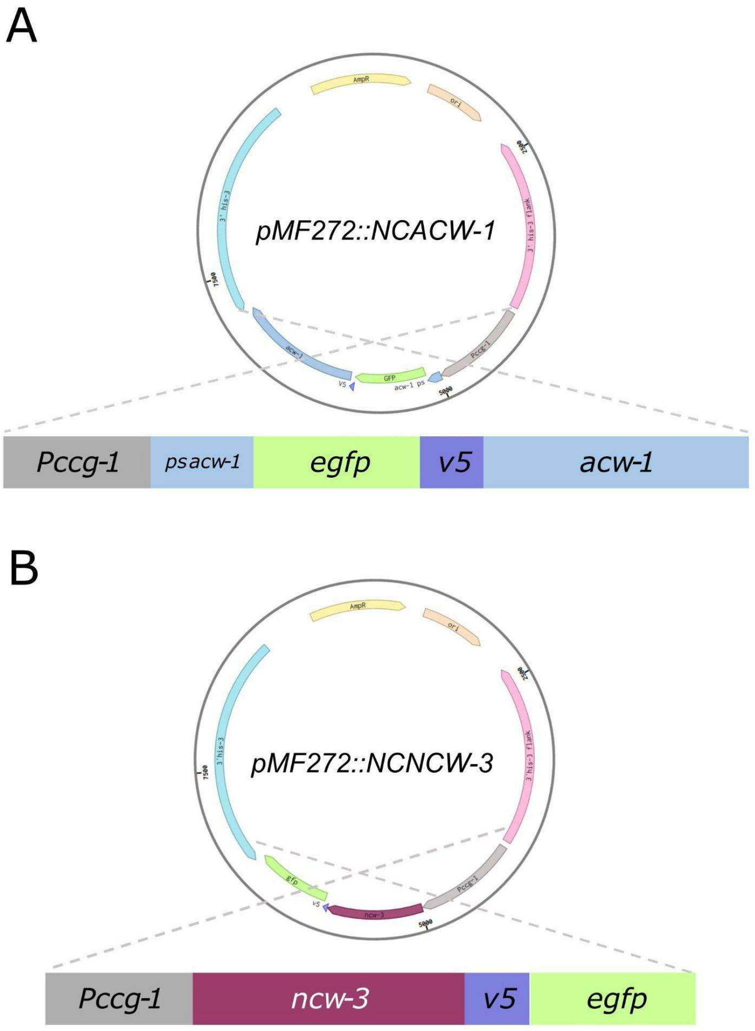


**Figure S5. Molecular design and construction of eGFP::V5::ACW-1 and NCW-3::V5::eGFP expression assemblies, *pMF272::NCACW-1* and *pMF272::NCNCW-3,* respectively.** Assemblies were placed under the control of the *ccg-1* promoter (*Pccg-1*), *egfp* was used as the fluorescent marker, and *v5* as epitope and spacer. *pMF272::NCACW-1* and *pMF272::NCNCW-3* were generated by Gibson assembly and cloned into the *pMF272* vector between 3' *his-3* and *his-3* downstream sequences to allow their homologous recombination into the *his-3* locus of the parental FGSC 9717 *N. crassa* strain (*Δmus-51::bar+::his$^{-3}$,* mat A). A) eGFP::V5::ACW-1 expression assembly, *pMF272::NCACW-1* (*Pccg-1::ps acw-1::egfp::v5::acw-1*). The native ACW-1 signal peptide sequence (*ps acw-1*) was used to direct the fusion to the cell surface. The *acw-1* gene sequence was fused in-frame to the N-terminal eGFP::V5 tag. B) NCW-3::V5::eGFP expression assembly, *pMF272::NCNCW-3* (*Pccg-1::ncw-3::v5::egfp*) used the native NCW-3 signal peptide sequence to direct the assembly to the cell surface, and the *ncw-3* gene sequence was fused in-frame to the C-terminal V5::eGFP tag. Correct assembly of *pMF272::NCACW-1* and *pMF272::NCNCW-3* was validated by sequencing.

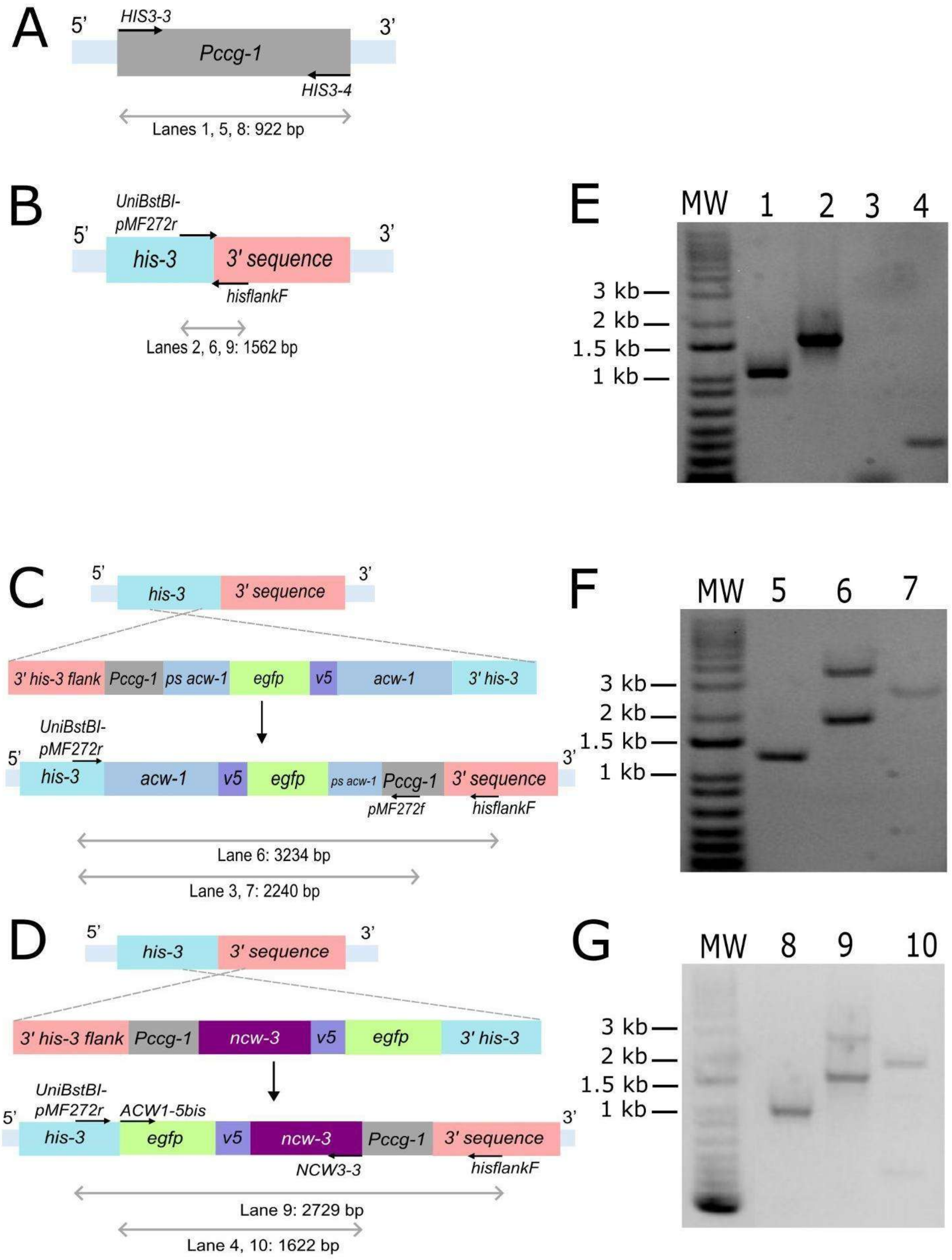


**Figure S6. Molecular characterization of constructed heterokaryotic *N. crassa* strains.** Assemblies *pMF272::NCACW-1* (*Pccg-1::ps acw-1::egfp::v5::acw-1*) and *pMF272::NCNCW-3* (*Pccg-1::ncw-3::v5::egfp*) were directed to the *his-3* locus of the parental FGSC 9717 *N. crassa* strain (*Δmus-51::bar+::his-3,* mat A) and integrated by homologous recombination to generate the Nc ACW-1 and Nc NCW-3 strains. A, E, F and G) PCR amplification of the loading control, *ccg-1* promoter (*Pccg-1*) with primers HIS3-3 and HIS3-4, from purified genomic DNA (gDNA) of Nc FGSC9717 parental strain (E, lane 1), Nc NCACW-1 (F, lane 5) and Nc NCNCW-3 (G, lane 8) strains. A band of expected size (922 bp) was obtained for all cases as the promoter was present in the three strains. B, C, D, E, F and G) PCR amplification with primers UniBstBI-pMF272r and hisflankF, that bind to the insertion region between 3' *his-3* and 3' downstream sequences, from purified gDNA of parental Nc FGSC 9717, Nc NCACW-1 and Nc NCNCW-3 strains. A band of the expected size (1562 bp), corresponding to the parental WT region in Nc FGSC strain (E, lane 2) appeared, which confirmed the WT sequence and absence of insertions. Bands around the same size (1562 bp) appeared for the Nc NCACW-1 (F, lane 6) and Nc NCNCW-3 (G, lane 9) strains, which corresponded to the untransformed parental WT region, while additional bands appeared about the expected sizes of the *pMF272::NCACW-1* (C, 3234 bp) for the Nc NCACW-1 (F, lane 6) and the *pMF272::NCNCW-3* (D, 2729 bp) for the Nc NCNCW-3 (G, lane 9). This confirmed the correct insertion of the assemblies and the heterokaryotic identities of the Nc NCACW-1 and Nc NCNCW-3 strains. C, E and F) PCR amplification of an inner region of the *pMF272::NCACW-1* assembly from gDNA of parental Nc FGSC9717 and Nc NCACW-1 strains, with primers UniBstBI-pMF272r (which binds to 3' *his-3*) and pMF272f (which binds to *Pccg-1*). As anticipated, there was no specific amplification for the parental Nc FGSC9717 strain (E, lane 3) due to the absence of integrated assemblies. In the Nc NCACW-1 strain (F, lane 7), a band of the expected size (2240 bp) appeared, thus confirming the presence of the *pMF272::NCACW-1* assembly from amplification of an inner region (C). D, E and G) PCR amplification of an inner region of the *pMF272::NCNCW-3* assembly from gDNA of parental Nc FGSC9717 and Nc NCNCW-3 strains, with primers ACW1-5Bis (which binds to *egfp*) and NCW3-3 (which binds to *ncw-3*). In the parental Nc FGSC 9717 strain (E, lane 4) a non-specific band appeared around 300 bp. In the Nc NCNCW-3 strain (G, lane 10), a band of the expected size (1622 bp) was amplified along with the non-specific band at 300 bp, confirming the presence of the *pMF272::NCNCW-3* from amplification of a region within the assembly (D).

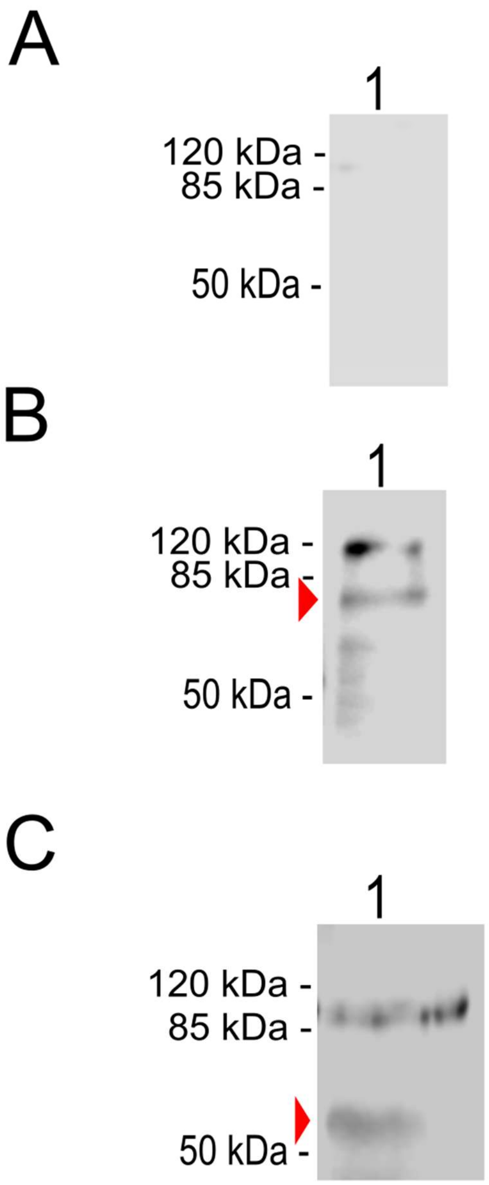


**Figure S7**. **Expression of eGFP-tagged ACW-1 and NCW-3 fusions.** *N. crassa* total homogenates were analyzed by Western Blot (Towbin et al., 1979) using V5 as the epitope to identify the expression of the eGFP-tagged ACW-1 and NCW-3 fusion proteins. For the parental Nc WT strain (A) no bands were detected except for a faint band between 120 and 85 kDa, calculated by relative mobility to be 104.4 kDa. This band also appeared for the Nc NCACW-1 (B) and Nc NCNCW-3 (C) strains, so they were considered background signals. For the Nc NCACW-1 strain (B), a band appeared below the 85 kDa molecular marker. The size of this band was calculated by relative mobility to be 84.9 kDa, which was similar to the expected size of the eGFP:V5:ACW-1 fusion (85.4 kDa: mature ACW-1, 57 kDa; V5 epitope, 1.4 kDa, and eGFP, 27 kDa). Lower size bands could represent partially digested versions of the eGFP-ACW-1 fusion. For the Nc NCNCW-3 strain (C), a band above the 50 kDa molecular marker was detected, calculated by relative mobility to be 54.7 kDa in size. This size was similar to the predicted size of the NCW-3:V5:eGFP fusion (52.0 kDa: NCW-3, 23.6 kDa; V5 epitope, 1.4 kDa, and eGFP, 27 kDa).

**Video S1. Z-stack reconstruction of *Neurospora crassa* NcNCACW-1 (*Pccg-1::ps acw-1::egfp::v5::acw-1*).** eGFP::V5::ACW-1 localized mainly around the septa of the distal regions of hyphae (white arrows) and the cell surface (yellow arrow). The video was edited in Fiji (Schindelin et al., 2012). Scale bar, 20 um.

**Video S2. Z-stack reconstruction of *Neurospora crassa* NcNCNCW-3 (*Pccg-1::ncw-3::v5::egfp*).** NCW-3::V5::eGFP was localized around the septa of the distal regions of hyphae (white arrows) and the cell surface (yellow arrow). The video was edited in Fiji (Schindelin et al., 2012). Scale bar, 20 um.
